# Long live the king: chromosome-level assembly of the lion (*Panthera leo*) using linked-read, Hi-C, and long read data

**DOI:** 10.1101/705483

**Authors:** Ellie E. Armstrong, Ryan W. Taylor, Danny E. Miller, Christopher Kaelin, Gregory Barsh, Elizabeth A. Hadly, Dmitri Petrov

## Abstract

The lion (*Panthera leo*) is one of the most popular and iconic feline species on the planet, yet in spite of its popularity, the last century has seen massive declines for lion populations worldwide. Genomic resources for endangered species represent an important way forward for the field of conservation, enabling high-resolution studies of demography, disease, and population dynamics. Here, we present a chromosome-level assembly for the captive African lion from the Exotic Feline Rescue Center as a resource for current and subsequent genetic work of the sole social species of the *Panthera* clade. Our assembly is composed of 10x Genomics Chromium data, Dovetail Hi-C, and Oxford Nanopore long-read data. Synteny is highly conserved between the lion, other *Panthera* genomes, and the domestic cat. We find variability in the length and levels of homozygosity across the genomes of the lion sequenced here and other previous published resequence data, indicating contrasting histories of recent and ancient small population sizes and/or inbreeding. Demographic analyses reveal similar histories across all individuals except the Asiatic lion, which shows a more rapid decline in population size. This high-quality genome will greatly aid in the continuing research and conservation efforts for the lion.

## Introduction

The lion (*Panthera leo*) is historically one of the most widespread carnivores on the planet, previously ranging from the tip of southern Africa, to the southern edge of North America [1,2]. However, over just the past 25 years, the African lion (*P. leo leo*) has lost more than half of its population, while the other recognized subspecies, the Asiatic lion (*P. leo persica*), has been reduced to fewer than 1,000 individuals, occupying little of their former range as a single population in the Gir forest, India. The remaining Asiatic lions are suspected to be suffering from reproductive declines due to inbreeding depression [3] and have been subject to several outbreaks of canine distemper virus (CDV;[4]).

Genetic markers have played a key role in studying the biogeography, history, and movement of lions for the past 50 years (e.g. [2,5,6], [7–10]). However, studies have been mostly limited to microsatellites with limited use of nuclear and mitochondrial genetic data (e.g. [11–17]). More recently, reduced representation sequencing has enabled genomic genotyping using the domestic cat or tiger as a reference [18]. Felid karyotypes are thought to be highly conserved [19,20], but studies have shown a reference mapping bias for estimation of statistics such as heterozygosity [21] and accurate allele calling [22], both of which are important for assessing population history.

The causes of the decline in lions are multifactorial. Lions have been hunted by humans for thousands of years, possibly first as a direct competitor and threat to our survival [23], for initiation rituals and rites of passage (e.g. [24–26]), to reduce predation of domesticated animals, and more recently for sport [27–30]. The recent rise of illegal trade in lion parts and illicit breeding practices has escalated over the past ten years, bringing hunting practices and international laws into the spotlight. In addition, several documentaries have exposed the lion breeding industry within South Africa, which uses fenced lions for ‘petting’, canned hunting experiences, and ultimately as skeletons for export, likely destined for Asian medicines [31]. Accurate and rapid genotyping could aid law enforcement to reveal whether the origins of trafficked goods are from wild or captive populations.

Moreover, rapid population decline has put lions at the forefront of the conservation debate over translocations and how best to manage populations. Many efforts to restore previous populations have focused on translocation lions within and between various South African lion populations (e.g. [32,33]. Information about local population adaptation, deleterious alleles, and potential inbreeding is lacking, which further complicates managed relocations. While increasing genetic diversity remains a widely accepted conservation goal, recent theoretical implications suggest consideration should be made when moving individuals from large heterozygous populations into small homozygous populations [34]. Genomic resources will aid immensely in these estimations and have already shown to be highly preferable to microsatellites or a reduced number of loci (see e.g. [35–37]).

To date, no *de novo* genome assembly for an African lion exists and only two individuals have been resequenced (Cho et al. (2013). A *de novo* assembly of an Asiatic lion was recently completed [38], but was limited to short-read technology, and was thus highly fragmented. Asiatic and African lions are currently regarded as separate subspecies [1,6,39], and we regard them as such for these analyses. Here, we present a high-quality, *de novo* genome assembly for the lion (*Panthera leo*), referred to as Panleo1.0. We use a combination of 10x Genomics linked-read technology, Dovetail Hi-C, and Oxford Nanopore long read sequencing to build a highly contiguous assembly. We verify the conserved synteny of the lion in comparison with the domestic cat assembly and also examine the demography and heterozygosity of the lion compared with other felids. It is our hope that this genome will enable a new generation of high quality genomic studies of the lion, in addition to comparative studies across *Felidae*.

## Results

### Genome assembly and continuity

The assembly generated with 10x Genomics Chromium technology yielded a consistent, high-quality starting assembly for the lion. In general, assembly statistics are improved when compared to previous assemblies generated using short-insert and mate-pair Illumina libraries, such as the tiger [56], cheetah [61], Amur leopard [62], lynx [63], and puma [64]. All these assemblies have improved their scaffold statistics through a variety of technologies, such as Pacbio, Bionano, Nanopore, or Hi-C (Supplementary Table S3; see publications above and DNAZoo). The lower contig scores are also emulated through a higher number of missing BUSCOs (Table 2, Supplementary Table S4). Although we were unable to compare it to the *de novo* assembly of the Asiatic lion from Mitra et al. (2019), they report a scaffold N50 of approximately 20kb, suggesting our assembly represents significant improvement.

**Table 1:** Assembly statistics for various assembly phases of the lion genome.

|  | **Lion (10x)** | **Lion (10x,Hi-C)** | **Panleo1.0 (10x, Hi-C, Nanopore)** |
| --- | --- | --- | --- |
| Number of scaffolds | 8,389 | 8,062 | 8,061 |
| Assembly size | 2.4 Gb | 2.4 Gb | 2.4 Gb |
| Mean scaffold size | 287 kb | 298 kb | 298 kb |
| Scaffold N50 | 10 Mb | 136 Mb | 136 Mb |
| Scaffold N90 | 20 Mb | 57 Mb | 57 Mb |
| Scaffold L50 | 66 | 8 | 8 |
| Scaffold L90 | 243 | 18 | 18 |
| Number of contigs | 36,266 | 36,266 | 22,589 |
| Mean contig size | 66 kb | 66 kb | 106 kb |
| Contig N50 | 161 kb | 161 kb | 312 kb |
| Contig N90 | 41,289 bp | 41,289 bp | 82,163 bp |
| Contig L50 | 4,376 | 4,376 | 2,262 |
| Contig L90 | 15,422 | 15,422 | 7,875 |
| # Ns | 22,827,550 | 22,860,250 | 18,222,230 |

**Table 2:**
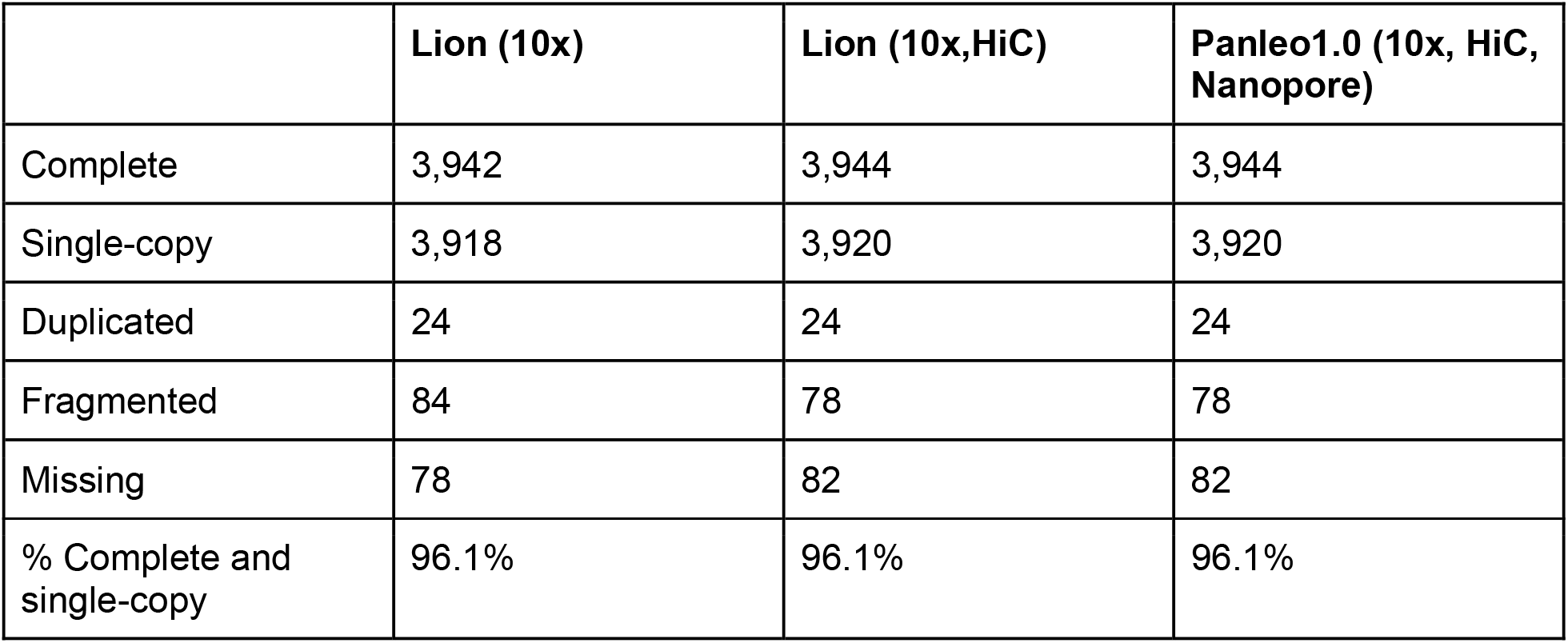
BUSCO scores for various assemblies.

### Using long sequencing reads to close gaps in draft genome assemblies

While the draft assemblies using either 10x alone or 10x + Dovetail Hi-C were of high-quality, they contained a number of gaps containing unknown sequence (Table 1). We therefore wondered if some of these gaps could be closed using long sequencing reads. Using a single Oxford Nanopore MinION flowcell we generated a total of 1,520,012 reads with an average read length of 6,528 bp, resulting in approximately 4x coverage of the *P. leo* genome. We then identified single reads which spanned gaps of any size and, for each gap, used MUSCLE and Cons to generate a consensus sequence of sequence spanning that gap (see methods). Using this approach we closed 26,403 gaps of 10, 100, or 400 bp with an average coverage of 3x per gap. We then identified split reads which spanned any gap 3kb or larger and again, for any instance in which multiple reads spanned a gap, pooled those reads and used MUSCLE and cons to generate a consensus sequence of the sequence spanning the gap. If only one read spanned the gap the raw sequence from that read was used to fill the gap. This approach resulted in the closing of 574 gaps of 3,000, 5,000, or 10,000 bp with an average coverage of 1x per gap. Overall, this approach closed 26,977 out of 42,635 gaps on 416 of the 8,061 scaffolds in the 10x + Dovetail assembly and reduced the overall size of the genome assembly by 1.6 million bp while increasing the mean contig size from 66 kb to 106 kb. Overall, this approach resulted in a substantial improvement on average contig size and associated statistics in the lion genome, but did not improve BUSCO scores for the genome. A detailed description of the gaps filled in using Nanopore can be found in Supplementary Table S2.

### Phylogenetics

To verify the phylogenetic relationships of the taxa using the *de novo* genomes, we constructed a phylogenetic tree using a maximum-likelihood approach. Consistent with recent phylogenetic analyses of the, we found that the lion, the leopard, and the tiger form a cluster representing *Panthera*, with the leopard and lion constituting sister species within the group [65,66]. The cheetah and puma comprise another cluster, with the lynx sitting outside this grouping [66]. The domestic cat is the most distantly related to all of the species tested here. Since we used protein files to infer the phylogenetic relationships, we found very high posterior probabilities across all the nodes (Figure 1).

**Figure 1:**
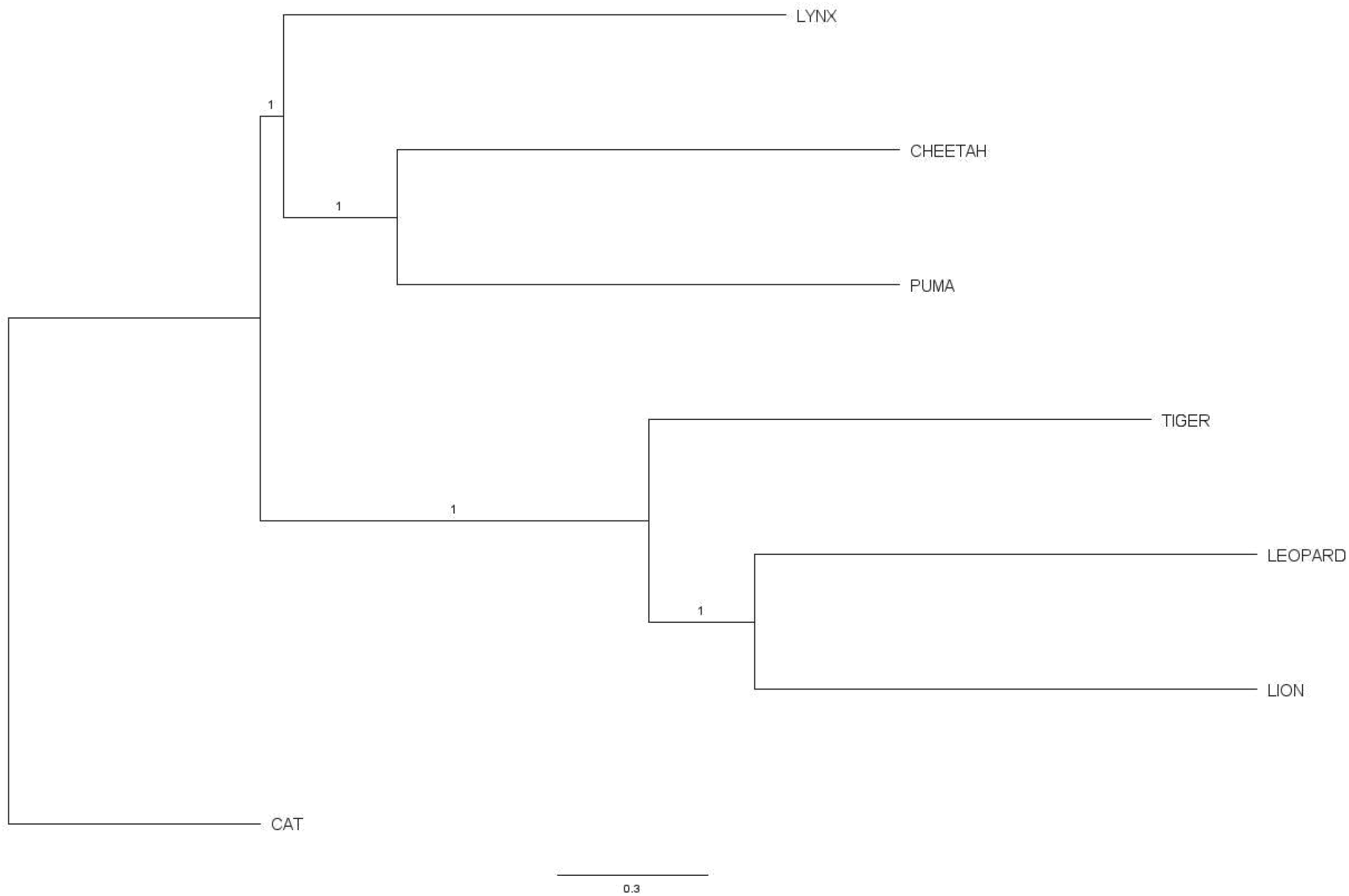
Phylogenetic reconstruction of *de novo* Felid genomes using RAxML and highly conserved genes from BUSCO mammalia_odb9 dataset. Node annotations indicate posterior probabilities.

### Repetitive Element and Gene Annotations

We generated statistics for repetitive elements in each genome using a pipeline which combines homology-based evidence and *de novo* repeat finding. On average, the continuity of the assembly did not greatly affect our ability to identify repeats (Table S5). Assemblies from each felid genome analyzed contained between 40.0%-42.5% repeats (Table S6). Alternatively, gene annotation results showed that more continuous assembles generate less annotated genes on average (Table S7, Table S8). Possibly, this indicates that more fragmented assemblies cause misidentifications of gene regions by automated annotation software.

### Synteny

We constructed genome synteny visualizations for chromosome-level assemblies of the domestic cat (*F. silvestris*: GCA_000181335), the lion (*P. leo*), and the tiger (*P. tigris*; [56], [57,58]). Each assembly was aligned to the domestic cat and the lion, in order to observe similarities and differences between the genomes. Consistent with expectation, we found very few large rearrangements in the karyotype across species (Figure 2, Supplementary Figures S1, S2).

**Figure 2:**
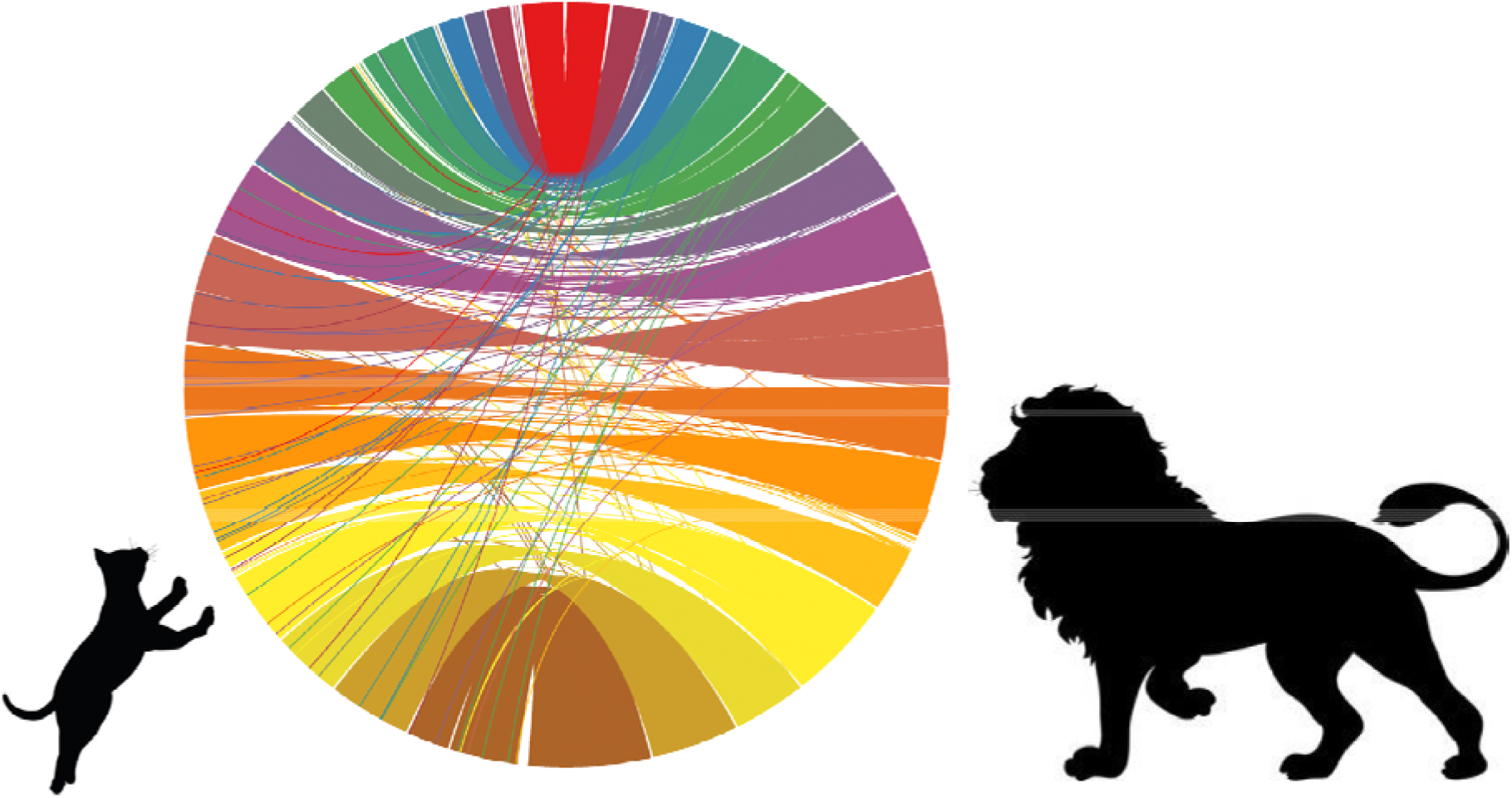
Circos plot of alignments between domestic cat (left) and lion (right) chromosomes. Colors represent different chromosomes with bottom chromosome (shown in dark brown) representing linkage group A1 of the domestic cat.

### Heterozygosity

We mapped raw Illumina reads to each respective species genome, as well as to the domestic cat assembly. We found that, on average, mapping to the domestic cat assembly resulted in lower heterozygosity calls and an average of 10% fewer reads successfully mapped (Table S10). However, this pattern was inconsistent and reversed for the Asiatic lion individual (Figure 3, Table S10). These results are supported by [21], who found that the reference used had some effect on heterozygosity inference, but little effect on the inference of population structure. In addition, we find that there is substantial variation in genome-wide heterozygosity estimates across the four lions that were tested (0.0012, 0.0008, 0.0007, and 0.00019, respectively). The two captive lions sequenced in Cho et al. (2013) may have been substantially inbred as part of their captive population, but no further details on the individuals are available.

**Figure 3:**
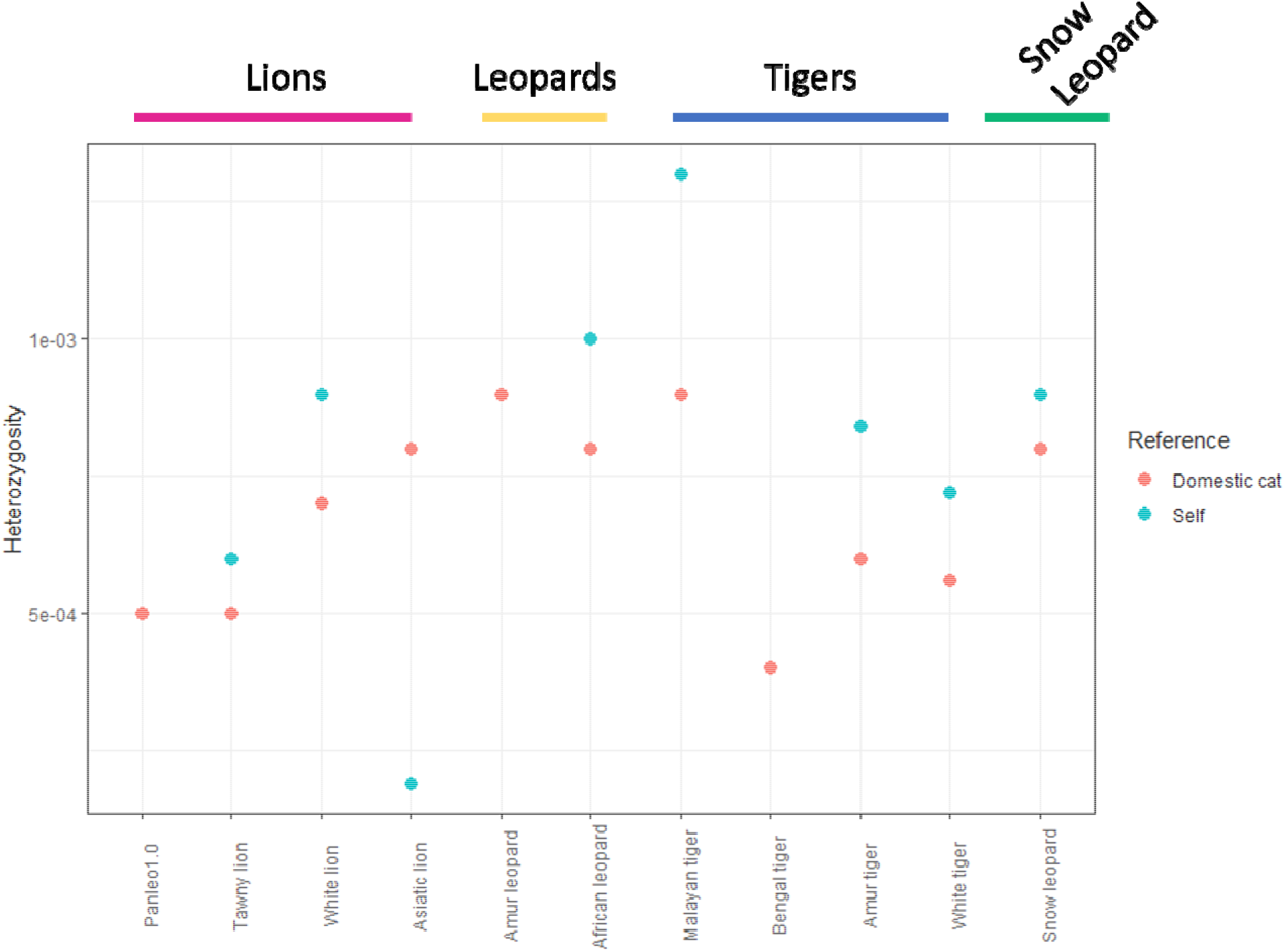
Average genome-wide heterozygosity of various Felids when mapped to a reference genome from their own species (blue) vs. when mapped to the domestic cat (red).

Because the assembly quality varied, we also performed two brief tests on *Panthera* datasets which have multiple reference genome assemblies (Table S9). We find that in general, fragmented assemblies do not seem to strongly influence heterozygosity calls, regardless of whether or not the individual in question represents the reference sequence.

### Runs of homozygosity

Using the mapped files created during the previous step, we investigated how runs of homozygosity were distributed across the four lion genomes. We found that there were a high proportion of short runs contained within the Asiatic lion genome (Figure S3), and to a lesser extent, the two previously published captive lion genome sequences from Cho et al. (2013). In general, heterozygosity was much lower genome-wide in the Asiatic individual (Figure 4), indicating that, along with showing signs of recent inbreeding, the population has likely been small for a long time (see [67]). Panleo1.0 (“Brooke”) had the highest overall heterozygosity, which may be a result of admixture that has occurred during unmonitored captive breeding which occurs outside of regulated zoos.

**Figure 4:**
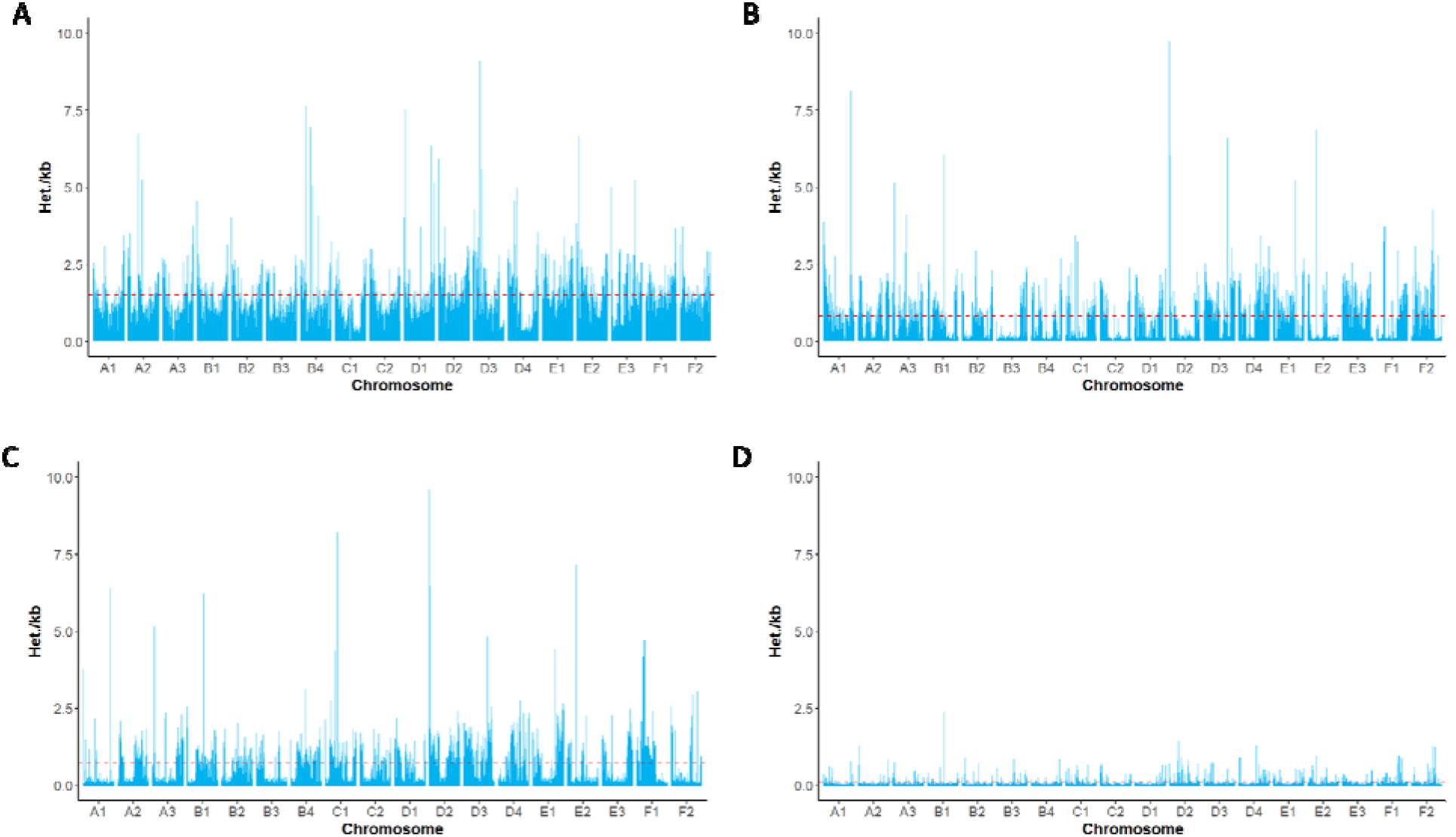
Genome-wide heterozygosity. Left panel shows heterozygosity genome-wide in nonoverlapping 1Mb bins. A: Lion from this study, “Brooke”, B: Tawny lion, Cho et al. (2013), C: White lion, Cho et al. (2013), D: Asiatic lion, Mitra et al. (2019). Red line represents the mean heterozygosity value genome-wide.

### Demographic history

PSMC analyses revealed similar demographic history of Panleo1.0 and the two genomes from Cho et al. (2013). These genomes show an initial, small decline approximately 1 million years ago (MYA), and a secondary more severe decline beginning approximately 500,000 years ago (Figure 5; Supplementary Figures S4, S5, S6). These trends are consistent with the fossil record which has revealed declines of large mammal populations during this time period, possibly due to Archaic human influence and/or climate changes (e.g. [68,69]). However, the Asiatic lion genome shows a more rapid decline over the past 100,000 years, but shows no consistent population size approximately 1MYA. It is possible that the low heterozygosity of the Asiatic lion was low enough to impede the inference of accurate historical N_E_ due to a distortion of the coalescent patterns across the genome. Corroborating these issues, other studies have shown variation between results in PSMC analyses within individuals of the same species and suggest that alternative coalescent methods should be used to confirm historical demographic trends [70]. Further, the genome sequenced here shows inconsistent signals in more recent history as the inference approaches the 100,000 year mark, which could be due to inflations of heterozygosity across the genome. Since the ancestral history of the individual sequenced here is unknown, it is possible that she is the result of crossings between multiple, distinct populations of lion which have variable histories.

**Figure 5:**
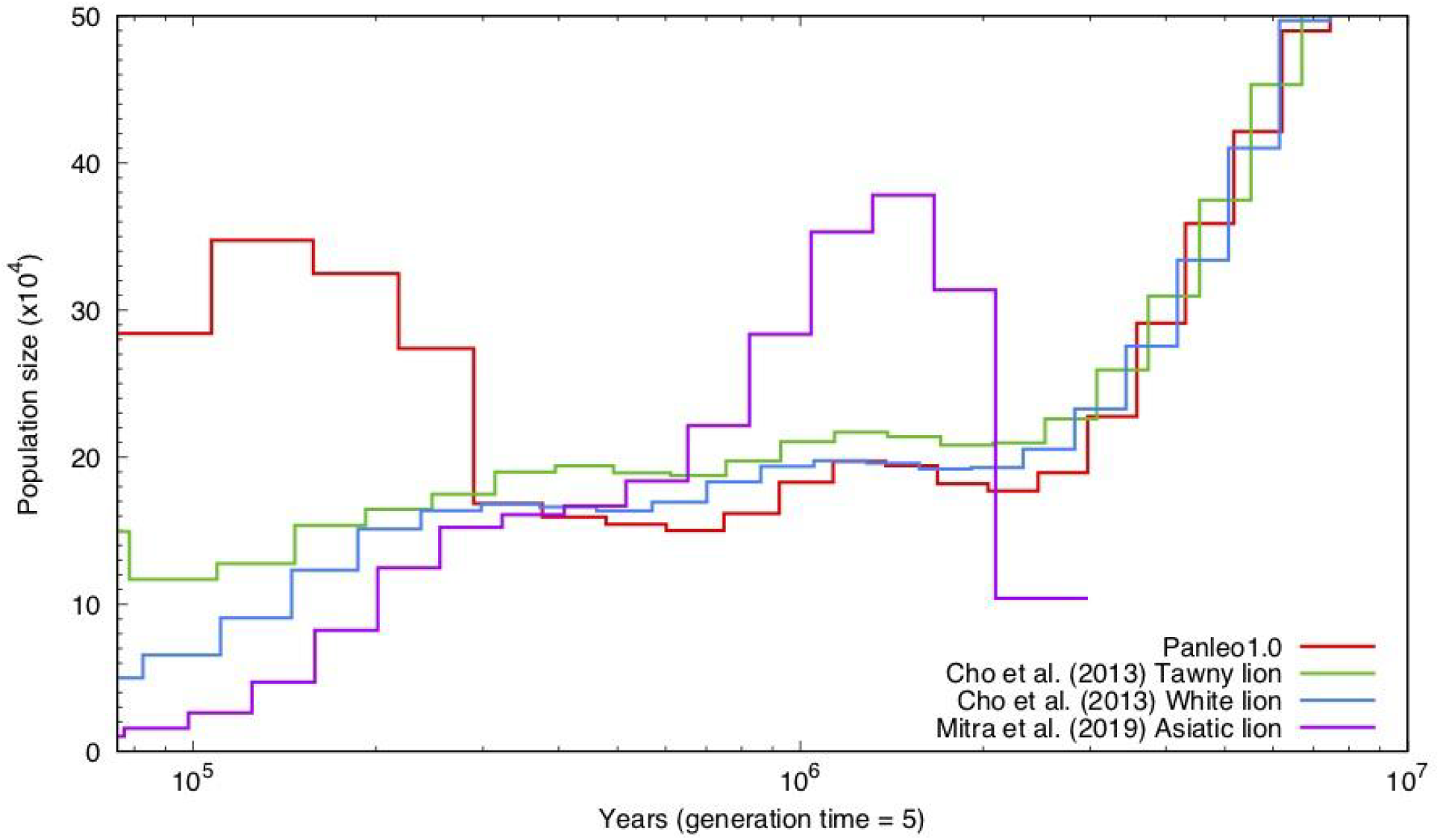
Demographic history of the lion as inferred by PSMC. Generation time used was 5 years and mutation rate applied was 0.1×10^−8^.

## Discussion

Long and linked-read genomic technologies such as 10x Genomics, Nanopore, and Hi-C allow rapid and economical *de novo* construction of high quality and highly contiguous genomes (e.g. [71]). Projects such as Genome 10k [72,73], i5k [74], DNAzoo (DNAzoo.org; [57,58]), and bird10k [75] aim to vastly improve our general understanding of the evolution of genomes, and both the origin and fate of diversity of life on earth. Such high quality assemblies will not only contribute to our understanding of the evolution of genomes, but also have practical applications in population genetics and conservation biology.

The chromosome-level *de novo* assembly of the lion genome presented here was constructed in three steps - 10x Genomics was used to create the base assembly and Dovetail Hi-C and Oxford Nanopore were used to improve contiguity. We show that each step results in substantial improvement to the genome, indicating that these methods are not redundant. At the same time, our data indicate that 10x and Hi-C alone are enough to approximate chromosomes in a typical mammal genome. Nanopore data, even with a small amount of very long reads, was enough to fill in many of the small gaps and ambiguous sequences across the genome.

The quality of this assembly allowed us to investigate the co-linearity of the genome compared to other felids and the importance of the reference sequence for estimating heterozygosity. As has been reported before [19,20], we find that the genomes of felids are largely co-linear and indicate that few large-scale chromosomal rearrangements have occurred across species. However, reference sequence bias can have substantial and unpredictable effect on estimating heterozygosity, possibly due to mismapping.

The variation of heterozygosity inference across the four lions tested here is further evidence that single genomes are not representative of the heterozygosity of a species or even the populations (captive or wild) from where they are derived. This assembly has also allowed us to compare fine-scale patterns of heterozygosity across the genome, where we find a substantial amount of variation between individuals. This contiguous genome will allow us to perform analyses on recent inbreeding and ROH in wild individuals across their range, how heterozygosity patterns differ between populations with different evolutionary histories, and how management decisions such as translocations and barriers to dispersal affect wild populations. Further, captive management of populations also stand to gain from genetic monitoring tools and, as we have shown here, individuals from zoos may harbor early signs of diversity loss and the accumulation of long runs of homozygosity. Even outside the nuanced case of the Asiatic lion, where dramatic population declines occurred prior to managers stepping in to monitor individuals, captive-bred populations often come from few founders with the addition of new individuals as available. If captive populations are truly meant to be a resource for conservation at large, more work must be done to understand the genetic implications of such scenarios.

Demographic analyses are also greatly aided by continuous sequence and rely on the inference of coalescence across the genome. As we detected a different historic demography for the Asiatic lion, it would be pertinent to examine how recent and rapid inbreeding affects the ability of these software to detect N_E_ over time. Further, examination of the patterns of diversity loss across wild individuals, especially populations which have been suggested to show signs of inbreeding (see the Ngorongoro crater lion population; [3,10,76]), will aid managers in decision making to ensure a future for existing lion populations.

This study will allow a surge forward in conservation efforts for the lion and enable studies across many facets of evolutionary biology, such as the highly conserved, yet phenotypic diversity of the genus *Panthera*. Undeniably, lion research has a historic legacy of collaboration across fields [77] and this genome will aid in future endeavors to prevent further loss of one of the world’s most iconic species. Most importantly, it will enable low-cost resequencing efforts to be completed, in addition to a wide range of other genetic studies, in order to further the conservation efforts of the lion.

## Methods

### Library Preparation and Sequencing

Whole blood samples were collected on two occasions during routine dental and medical procedures on an adult female lion (“Brooke”) from the Exotic Feline Rescue Center in 2017. Blood was collected in EDTA tubes, briefly held at −20C before being shipped overnight to Stanford and subsequently frozen at −80C. Approximately 1mL of whole blood was used for 10x Genomics Chromium library preparation and sequencing at HudsonAlpha in Huntsville, Alabama. This library was sequenced on an Illumina HiSeq X Ten. An additional 1mL was then sent to Dovetail Genomics in Santa Cruz, California for HiC library preparation and subsequent sequencing on the Illumina HiSeq X platform.

DNA for Nanopore sequencing was extracted from three 500uL aliquots of whole blood using the Quiagen DNeasy kit following the manufacturer’s instructions. DNA was eluted into 50uL and then concentrated to approximately 25ng/μL using a Zymo DNA Clean and Concentrator Kit. The final elution volume after concentrating was approximately 50μL. Libraries for Nanopore sequencing were prepared using a 1D genomic ligation kit (SQK-LSK108) following the manufacturer’s instructions with the following modifications: da-tailing and FFPE repair steps were combined by using 46.5μL of input DNA, 0.5μL NAD+, 3.5μL Ultra II EndPrep buffer and FFPE DNA repair buffer, and 3.0μL of Ultra II EndPrep Enzyme and FFPE Repair Mix, for a total reaction volume of 60μL. Subsequent thermocycler conditions were altered to 60 minutes at 20C and 30 minutes at 65C. The remainder of the protocol was performed according to the manufacturer’s instructions. 15μl of the resulting library was loaded onto a MinION with a R9.4.1 flowcell and run for 48 hours using MinKNOW version 2.0. Fastq files were generated from raw Nanopore data using Albacore version 2.3.1. Pass and fail reads were combined for a total of 1,520,012 reads with an average read length of 6,528 bp, with 336,792 of these reads greater than 10kb, and a longest read length of 62,463 bp.

### Genome Assembly

The 10x reads were assembled using Supernova version 1.2.1 with standard settings [40]. A single version of the genome was output using the ‘--pseudohap 1’ flag. This assembly was then provided to the HiRise software [41] as the starting assembly. HiRise was performed by Dovetail Genomics (Santa Cruz, CA) and the resulting assembly returned to us.

### Using long sequencing reads to close assembly gaps

Long sequencing reads generated by Nanopore sequencing were used to close gaps in the final 10x + Dovetail assembly. First, all Nanopore reads were mapped to the 10x + Dovetail Hi-C assembly using BWA [42] with the ont2d option (flags: -k14 -W20 -r10 -A1 -B1 -O1 -E1 - L0). Gaps were then closed using one of two methods, depending on whether single reads mapped to both sides of the gap.

We first identified single reads that had not been split by the aligner that mapped to at least 50 bp of sequence on either side of a gap in the 10x + Dovetail assembly and found 110,939 reads meeting this criteria. The sequence spanning the gap plus 50 bp on either side was extracted from the read and combined with other reads spanning the same gap into a single fasta file. To improve the quality of the alignment, 50 bp of sequence from either side of the gap from the reference genome was added to the fasta file. MUSCLE version 3.8.31 [43] was used, with default settings, to generate a multiple sequence alignment using all input sequences for each gap. Cons version 6.5.7.0 [44] was used to create a consensus sequence from the multiple alignment generated by MUSCLE. For each consensus sequence all N’s were removed.

Gaps not closed by single reads were then filtered and instances in which a single read was split and mapped to either side of a gap were identified, revealing 841 reads meeting this criteria. The sequence that spanned the gap but was not mapped was isolated and the 50 bp of sequence from the reference genome was added to either side of the unmapped sequence in a fasta file containing all gaps. In those instances where more than one split read spanned a gap MUSCLE was used to generate a multiple sequence alignment and Cons was then used to create a consensus sequence. Gaps in the reference genome were then replaced with the new consensus sequence.

### Quality Assessment

In order to assess the continuity of each genome assembly, we first ran scripts from Assemblathon 2, which gives a detailed view of the contig and scaffold statistics of each genome [45]. We then ran BUSCOv3 [46] in order to assess the conserved gene completeness across the genomes. We queried the genomes with the mammalian_odb9 dataset (4104 genes in total). We ran all three versions of the genome assembled here (10x, 10x + Hi-C, and 10x + Hi-C + Nanopore). The final version of the assembly (10x + Hi-C + Nanopore) is what we refer to as Panleo1.0.

We also used the genes queried by BUSCOv3 in order to infer phylogenetic relationships among the felids with assembled genomes (see Table S1 for details). We first extracted all the genes in the mammalia_odb9 dataset produced for each genome by each independent BUSCO run. These protein sequences were then aligned using MAAFT ([47]; flags ‘--genafpair’ and ‘--maxiterate 10000’). We then used RAxML [48] to build phylogenies for each of the genes. We used flags ‘-f a’, ‘-m PROTGAMMAAUTO’, ‘-p 12345’, ‘-x 12345’, ‘-# 100’, which applied a rapid bootstrap analysis (100 bootstraps) with a GAMMA model for rate heterogeneity. Flags ‘-p’ and ‘-x’ set the random seeds. We subsequently used the ‘bestTree’ for each gene and ran ASTRAL [49] on the resulting trees (3,439 trees total) to output the best tree under a maximum-likelihood framework.

### Repeat Masking

We identified repetitive regions in the genomes in order to perform repeat analysis and to prepare the genomes for annotation. Repeat annotation was accomplished using homology-based and *ab-initio* prediction approaches. We used the felid RepBase (http://www.girinst.org/repbase/; [50]) repeat database for the homology-based annotation within RepeatMasker (http://www.repeatmasker.org; [51]). The RepeatMasker setting -gccalc was used to infer GC content for each contig separately to improve the repeat annotation. We then performed *ab-initio* repeat finding using RepeatModeler (http://repeatmasker.org/RepeatModeler.html; [52]). RepeatModeler does not require previously assembled repeat databases and identifies repeats in the genome using statistical models. We performed two rounds of repeat masking for each genome. We first hard masked using the ‘-a’ option and ‘-gccalc’ in order to calculate repeat statistics for each genome. We subsequently using the ‘-nolow’ option for soft-masking, which converts regions of the genome to lower case, but does not entirely remove them. The soft-masked genome was used in subsequent genome annotation steps.

### Annotation

Gene annotation was performed with the Maker3 annotation pipeline using protein homology evidence from the felid, human, and mouse UniProt databases. Gene prediction was performed with the Augustus Human model. We calculated annotation statistics on the final ‘gff’ file using jcvi tools ‘-stats’ option [53].

### Synteny

We identified scaffolds potentially corresponding to chromosomes and any syntenic rearrangements between species. To do this we used the LAST aligner [54] to align the 20 largest scaffolds from each assembly to the linkage groups established by felcat9 (NCBI: GCA_000181335). We first created an index of each genome using the ‘lastdb’ function with flags ‘-P0’, ‘-uNEAR’, and ‘-R01’. We then determined substitutions and gap frequencies using the ‘last-train’ algorithm with flags ‘-P0’, ‘--revsym’, ‘--matsym’, ‘--gapsym’, ‘-E0.05’, and ‘-C2’. We then produced many-to-one alignments using ‘lastal’ with flags ‘-m50’, ‘-E0.05’, ‘-C2’ and the algorithm ‘last-split’ with flag ‘-m1’. Many-to-one alignments were filtered down to one-to-one alignments with ‘maf-swap’ and ‘last-split’ with flag ‘-m1’. Simple sequence alignments were discarded using ‘last-postmask’ and the output converted to tabular format using ‘maf-convert -n tab’. Alignments were then visualized using the CIRCA software (http://omgenomics.com/circa) and mismap statistics calculated.. We did not visualize any alignments that had an error probability greater than 1×10^-5. We additionally did not plot the sex chromosomes due to excessive repetitive regions and differences between the sexes of the animals that we used.

### Heterozygosity

Raw illumina reads from each species were mapped to the domestic cat genome (NCBI: GCA_000181335) and the reference genome for each respective species using BWA-MEM [42]. Observed heterozygosity was calculated using ANGSDv0.922 [55]. We first estimated the SFS for single samples using the options ‘-dosaf 1’, ‘-gl 1’, ‘-anc’, ‘-ref’, ‘-C 50’, ‘-minQ 20’, -’fold 1’, and ‘-minmapq 30’ (where ‘-anc’ and ‘-ref’ were used to specify the genome it was mapped to). Subsequently, we ran ‘realSFS’ and then calculated the heterozygosity as the second value in the site frequency spectrum.

To control for possible differences in heterozygosity due to mapping or assembly quality, we also performed the same analyses on genome assemblies of different qualities for the lion (*P.leo*; this study, 10x and 10x + HiC + Nanopore), and the tiger (*P. tigris*; [56], [57,58], [59]).

### Runs of homozygosity

Mapped sequences subsequently were used to infer runs of homozygosity across the genome. We used the ‘mafs’ output files from an additional run using ANGSD by adding the filters ‘-GL 1’, ‘-doMaf 2’, ‘-SNP_pval 1e-6’, ‘-doMajorMinor 1’, ‘-only_proper_pairs 0’, and ‘-minQ 15’. This run outputs a file that contains the positions of heterozygous sites across the genome. We counted the number of heterozygous sites in 1Mb bins across each scaffold and computed 1) the number of heterozygous sites in each bin and 2) the frequency of bins containing the number of heterozygous sites per kilobase. We then visualized this across the chromosomes as a proxy for runs of homozygosity in the genome, in addition to the length of the runs.

### Demographic history

We removed small scaffolds (cutoff of L90 value for each genome) and those which aligned to the cat X chromosome from the lion (see LAST mapping section above). The remaining scaffolds were used for PSMC. Reads were mapped to the remaining scaffolds using BWA-MEM and the consensus sequence called using SAMtools mpileup [60], BCFtools call, and vcfutils ‘vcf2fastq’. Minimum depth cutoffs of 10 and maximum depth cutoffs of 100 were applied to all genomes using vcfutils.In order to visualize the PSMC graphs, we applied a mutation rates of 0.9e-09 [56] and a generation time of five years for the lion. We compared these inferences with those from two previously resequenced lions [56] and the Asiatic lion [38].

## Acknowledgements

We thank J. Taft, J. Herrberg, R. Rizzo and the rest of the staff and volunteers of Exotic Feline Rescue Center, Center Point, Indiana, for providing access to samples from Brooke the lion. We thank K. Reeves and B. Nimmo of Tigers in America for coordination of samples and their continuing support of this project and the Stanford Program for Conservation Genomics. We also thank the University of Illinois College of Veterinary Medicine Urbana-Champagne, the Peter Emily Foundation, and Dr. G. Weber-Reid for assistance with sample transfers and access. We also thank K. Panchenko for assistance writing assembly quality scripts. We thank M. Daly of Dovetail Genomics for assistance with the Dovetail submission and processing. We thank E. Ebel, G. Battu of HudsonAlpha, for assistance with lab work and sequencing. A special thanks to T. Yokoyama for assistance with Nanopore sequencing and preparation and C. Yakym for assistance with making plots. Thanks to K. Solari and S. Morgan for helpful comments on the manuscript. Animal silhouettes used are from shutterstock.com (IDs:94265464 and 279191219). We fondly remember Brooke the lion, who passed away July 2019.

## Author Contributions

E.E.A. conceived of the study, performed genome assembly and analysis. R.W.T. performed genome annotation and analysis. D.E.M. performed scaffolding using Nanopore data. G.B. and C.K. assisted with genome sequencing and assembly. E.E.A., R.W.T, D.E.M., E.A.H., and D.P. wrote the manuscript. All authors approved the manuscript.

## Data Availability

Data and scripts used to produce these analyses are available upon request from the authors. Print will be updated with NCBI information when available.

## Supplementary Tables

**Table S1:**
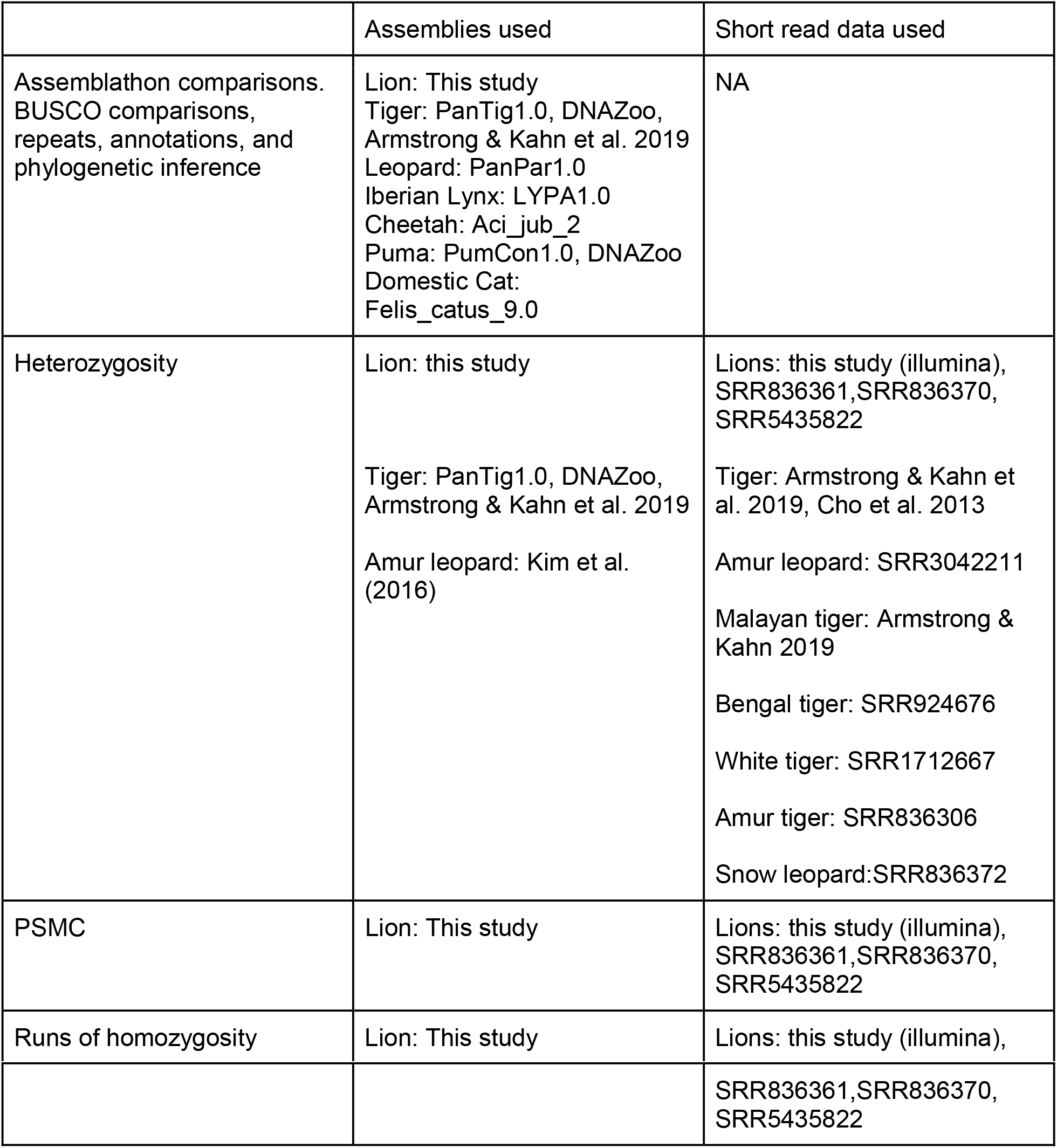
Summary of data sources used for analyses

**Table S2:**
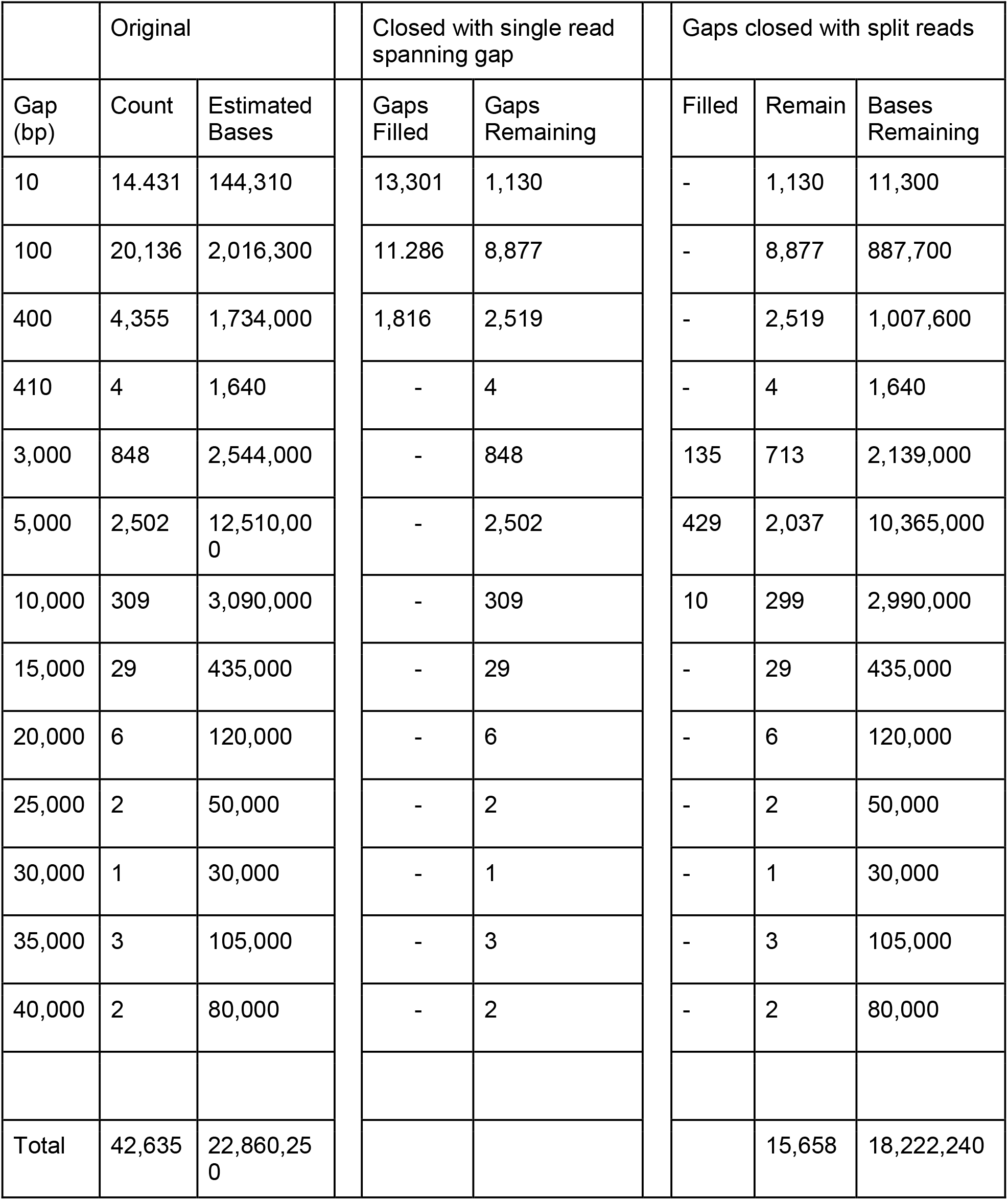
Details of genome assembly fill in with Oxford Nanopore data.

**Table S3:** Comparative assembly statistics from Assemblathon from published Felid genomes.

| Number of scaffolds | Lion | Tiger | Leopard | Iberian Lynx | Cheetah | Puma | Domestic Cat |
| --- | --- | --- | --- | --- | --- | --- | --- |
| Assembly size | 2.4 | 2.4 | 2.6 | 2.4 | 2.4 | 2.4 | 2.5 |
| Number of Scaffolds | 8,061 | 10,077 | 50,377 | 41,700 | 14,383 | 2,175 | 4,508 |
| Mean scaffold size | 298 kb | 241 kb | 51 kb | 58 kb | 166 kb | 1 Mb | 559 kb |
| Scaffold N50 | 136 Mb | 21 Mb | 21 Mb | 1.5 Mb | 67 Mb | 100 Mb | 150 Mb |
| Scaffold L90 | 18 | 109 | 135 | 2,019 | 780 | 26 | 16 |
| Number of contigs | 22,589 | 36,444 | 162,036 | 86,807 | 140,109 | 157,566 | 4,815 |
| Contig N50 | 312 kb | 186 kb | 41 kb | 107 kb | 47 kb | 32 kb | 514.3 kb |
| Mean contig size | 106 kb | 65 kb | 15 kb | 27 kb | 59 kb | 15 kb | 42 Mb |

**Table S4:**
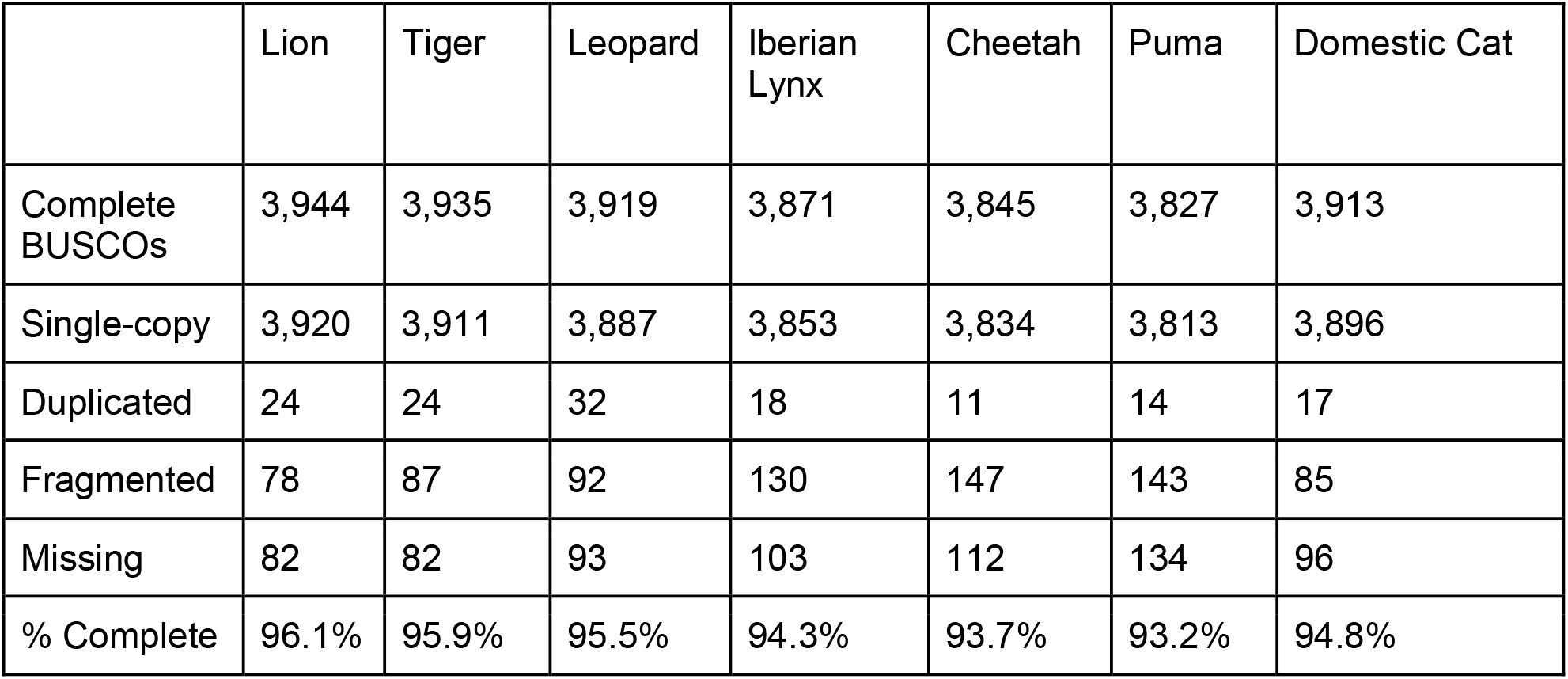
Comparative BUSCO scores between published Felid assemblies.

**Table S5:** Repeat statistics for *de novo* genome assemblies.

| Assembly | LINE | SINE | LTR | DNA | Unclassified | SmRNA | Others | Total (%) |
| --- | --- | --- | --- | --- | --- | --- | --- | --- |
| Lion (10x) | 889,658 | 1,468,544 | 315,693 | 342,778 | 13,913 | 1,099,269 | 1,018,633 | 42.30% |
| Lion (10x, Hi-C) | 891,625 | 1,469,034 | 315,965 | 342,983 | 13,524 | 1,099,584 | 1,018,250 | 42.33% |
| Lion (10x, Hi-C, Nanopore) | 890,292 | 1,469,373 | 317,748 | 338,708 | 17,853 | 1,099,878 | 1,016,325 | 42.45% |

**Table S6:** Repeat statistics for various *Panthera* assemblies and the domestic cat (felcat9).

| Species | LINE | SINE | LTR | DNA | Unclassified | SmRNA | Others | Total(%) |
| --- | --- | --- | --- | --- | --- | --- | --- | --- |
| Lion | 890,292 | 1,469,373 | 317,748 | 338,708 | 17,853 | 1,099,878 | 1,016,325 | 42.45 |
| Leopard | 944,218 | 1,564,679 | 330,933 | 354,148 | 14,427 | 1,184,163 | 1,090,064 | 40.97 |
| Tiger | 892,411 | 1,467,533 | 318,598 | 343,750 | 16,369 | 1,097,200 | 1,008,596 | 42.20 |
| Jaguar | 905,379 | 1,492,875 | 321,547 | 345,501 | 15,402 | 1,121,374 | 1,045,682 | 42.43 |
| Domestic cat | 891,936 | 1,555,928 | 318,675 | 346,001 | 19,497 | 1,183,922 | 1,036,956 | 42.04 |

**Table S7:**
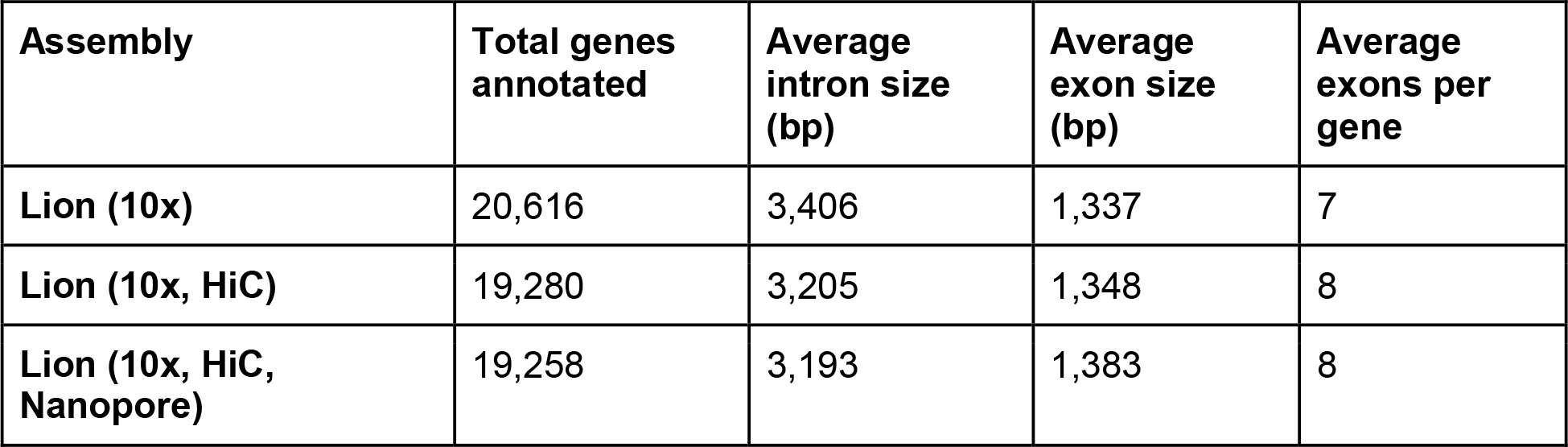
Annotation statistics for *de novo* assemblies from the jcvi program.

**Table S8:** Annotation statistics for *Panthera* genome assemblies and the domestic cat (felcat9) using jcvi.

| Assembly | Total genes annotated | Average intron size (bp) | Average exon size (bp) | Average exons per gene |
| --- | --- | --- | --- | --- |
| Lion (Panleo1.0) | 19,258 | 3,193 | 1,383 | 8 |
| Tiger (Cho et al. 2013) | 20,222 | 3,352 | 1,339 | 7 |
| Tiger (Armstrong & Kahn et al. 2019) | 20,261 | 3,045 | 1,401 | 8 |
| Tiger 3 (Cho et al. 2013, DNAZoo) | 17,953 | 3,447 | 1,281 | 7 |
| Leopard (Kim et al. 2016) | 20,864 | 3,476 | 1,335 | 7 |
| Leopard (Kim et al. 2016, DNAZoo) | 19,750 | 3,348 | 1,360 | 8 |
| Domestic Cat | 17,111 | 3,197 | 1,344 | 8 |

**Table S9:** Observed heterozygosity statistics from various assembly versions of the lion and tiger.

| Assembly | Heterozygosity | Number reads mapped | Avg. depth |
| --- | --- | --- | --- |
| Tiger (Cho et al.2013) | 0.0010 | 566,730,100 | 35.6 |
| Tiger (Cho et al. 2013, DNA zoo) | 0.0010 | 566,730,225 | 35.6 |
| Tiger (Armstrong & Kahn et al. 2019) | 0.0010 | 570,020,595 | 35.1 |
| Lion (10x-this study) | 0.0013 | 869,811,746 | 46.0 |
| Lion (Dovetail, 10x, nanopore-this study) | 0.0013 | 869,867,776 | 46.0 |

**Table S10:** Heterozygosity (observed) from various *Panthera* individuals when mapped to their own genome compared to when mapped to the domestic cat.

| Individual | Heterozygosity<br>(mapped to self) | Number reads mapped | Avg. depth | Heterozygosity<br>(mapped to felcat9) | Number reads mapped | Avg. depth |
| --- | --- | --- | --- | --- | --- | --- |
| Panleo1.0 | 0.00120 | 869,867,776 | 46.0 | 0.0009 | 866,802,126 | 45.5 |
| Tawny lion<br>(Cho et al. 2013) | 0.00084 | 801,148,358 | 30.1 | 0.0006 | 797,619,906 | 29.8 |
| White lion<br>(Cho et al. 2013) | 0.00072 | 1,416,024,084 | 53.2 | 0.0056 | 1,409,255,185 | 52.5 |
| Asiatic lion | 0.00019 | 516,572,489 | 23.4 | 0.0008 | 542,324,542 | 22.4 |
| Amur leopard<br>(Kim et al. 2016) | 0.0006 | 1,829,792,054 | 37.0 | 0.0005 | 908,350,242 | 37.3 |
| African leopard<br>(Kim et al. 2016) | 0.0005 | 370,623,557 | 15.0 | 0.0005 | 367,446,294 | 15.3 |
| Malayan tiger<br>(Armstrong et al. 2019) | 0.0010 | 570,020,595 | 35.1 | 0.0008 | 568,170,003 | 34.8 |
| Bengal tiger<br>(Cho et al. 2013) | 0.0009 | 839,375,249 | 31.9 | 0.0009 | 834,272,013 | 31.7 |
| Amur tiger<br>(Cho et al. 2013) | 0.0009 | 727,692,050 | 29.8 | 0.0007 | 721,809,185 | 29.5 |
| White tiger<br>(Cho et al. 2013) | 0.0009 | 727,134,421 | 27.8 | 0.0008 | 723,139,361 | 27.6 |
| Snow leopard<br>(Cho et al. 2013) | N/A | N/A | N/A | 0.0004 | 823,300,505 | 30.2 |

## Supplementary Figures

**Figure S1:**
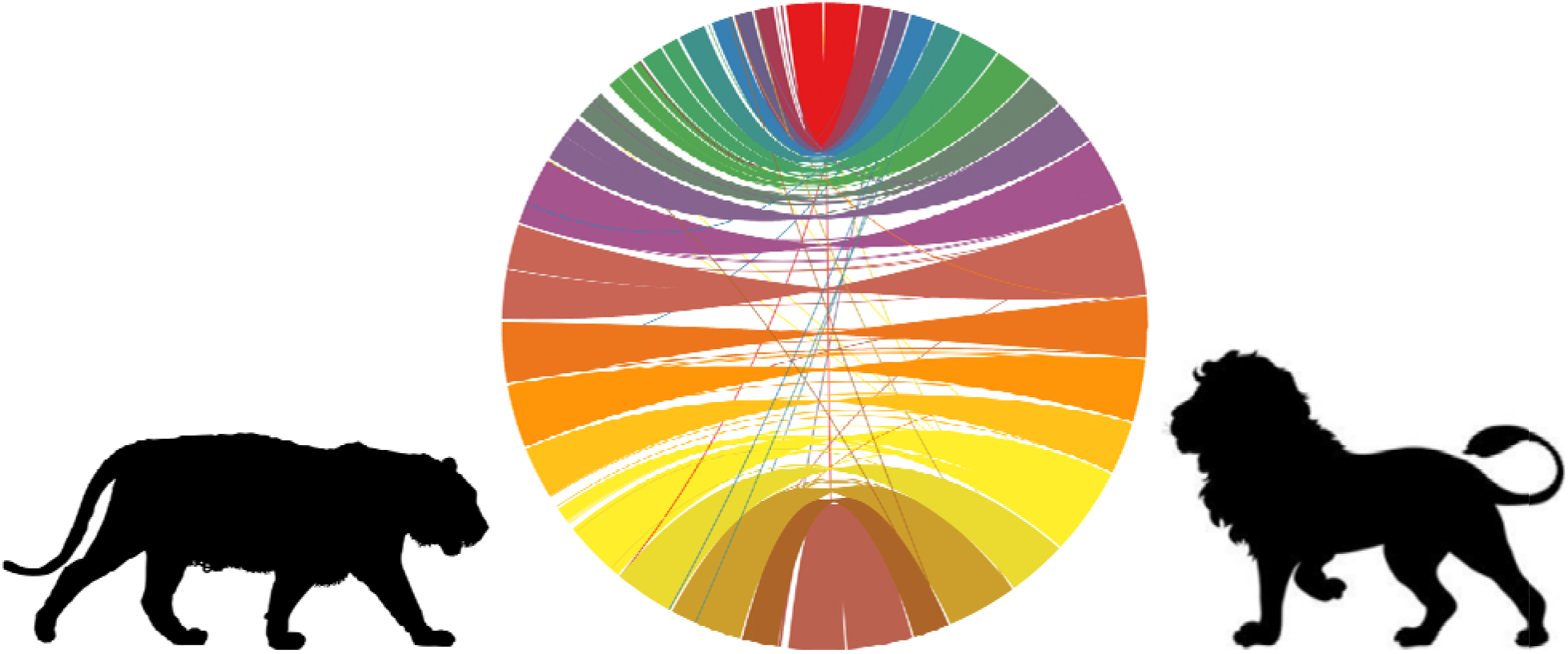
Circos plot of alignments between tiger (left) and lion (right) chromosomes. Colors represent different chromosomes with bottom chromosome (shown in dark brown) representing A1.

**Figure S2:**
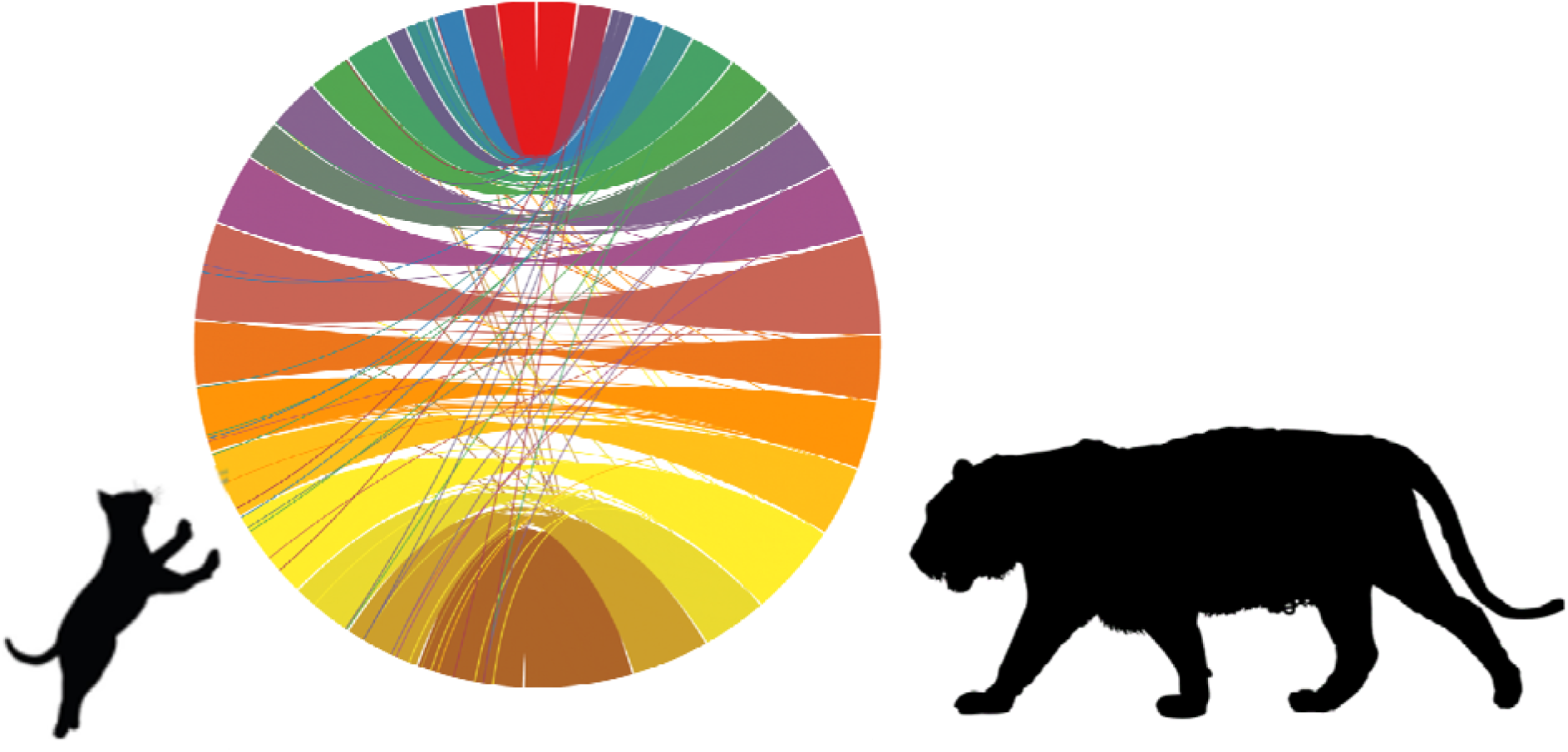
Circos plot of alignments between tiger (right) and domestic cat (left) chromosomes. Colors represent different chromosomes with bottom chromosome (shown in dark brown) representing A1.

**Figure S3:**
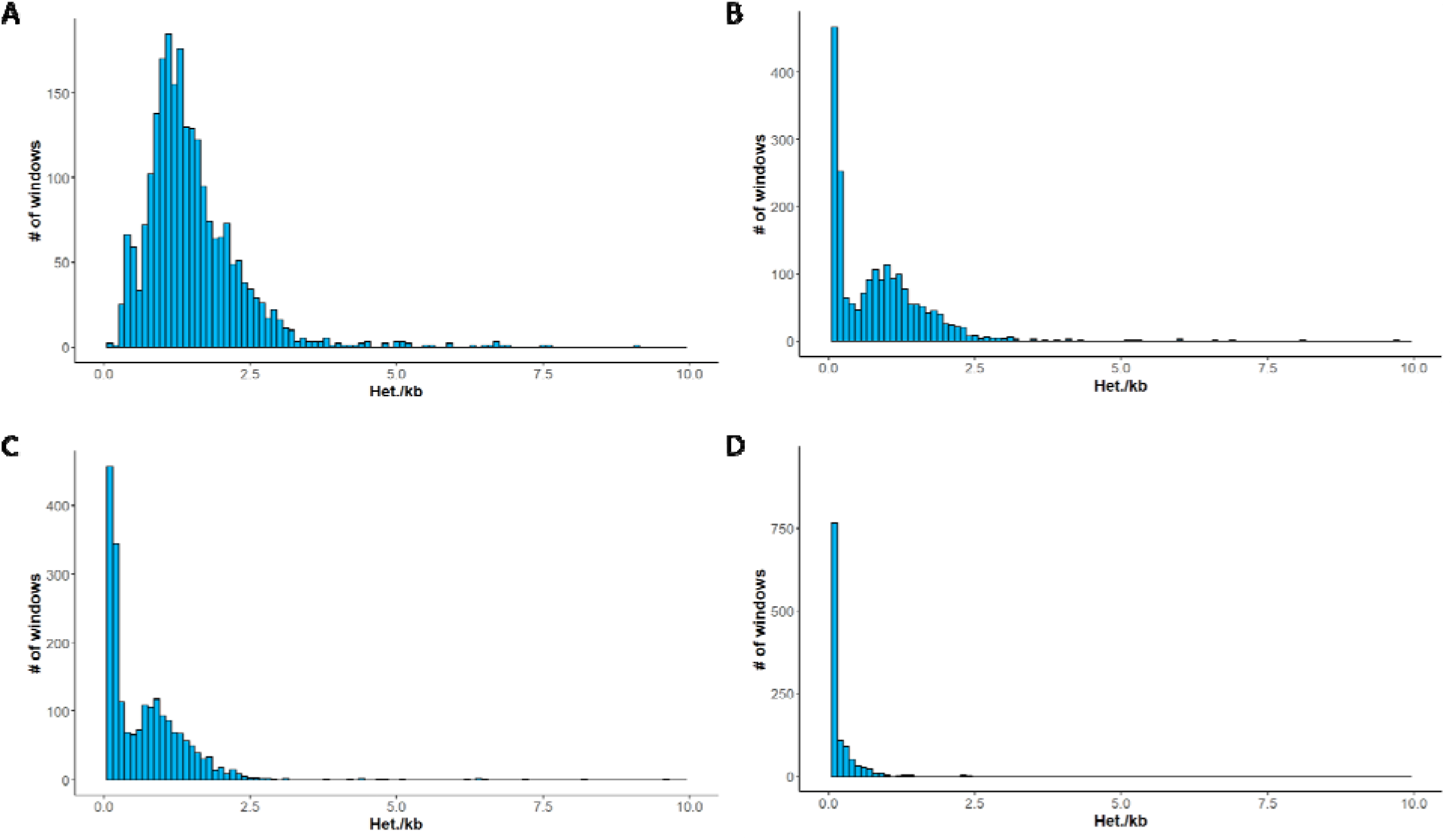
Histograms of per window heterozygosity. Graphs skewed more left represent individuals with more windows having lower heterozygosity on average. A: Lion from this study, “Brooke”, B: Tawny lion, Cho et al. (2013), C: White lion, Cho et al. (2013), D: Asiatic lion, Mitra et al. (2019).

**Figure S4:**
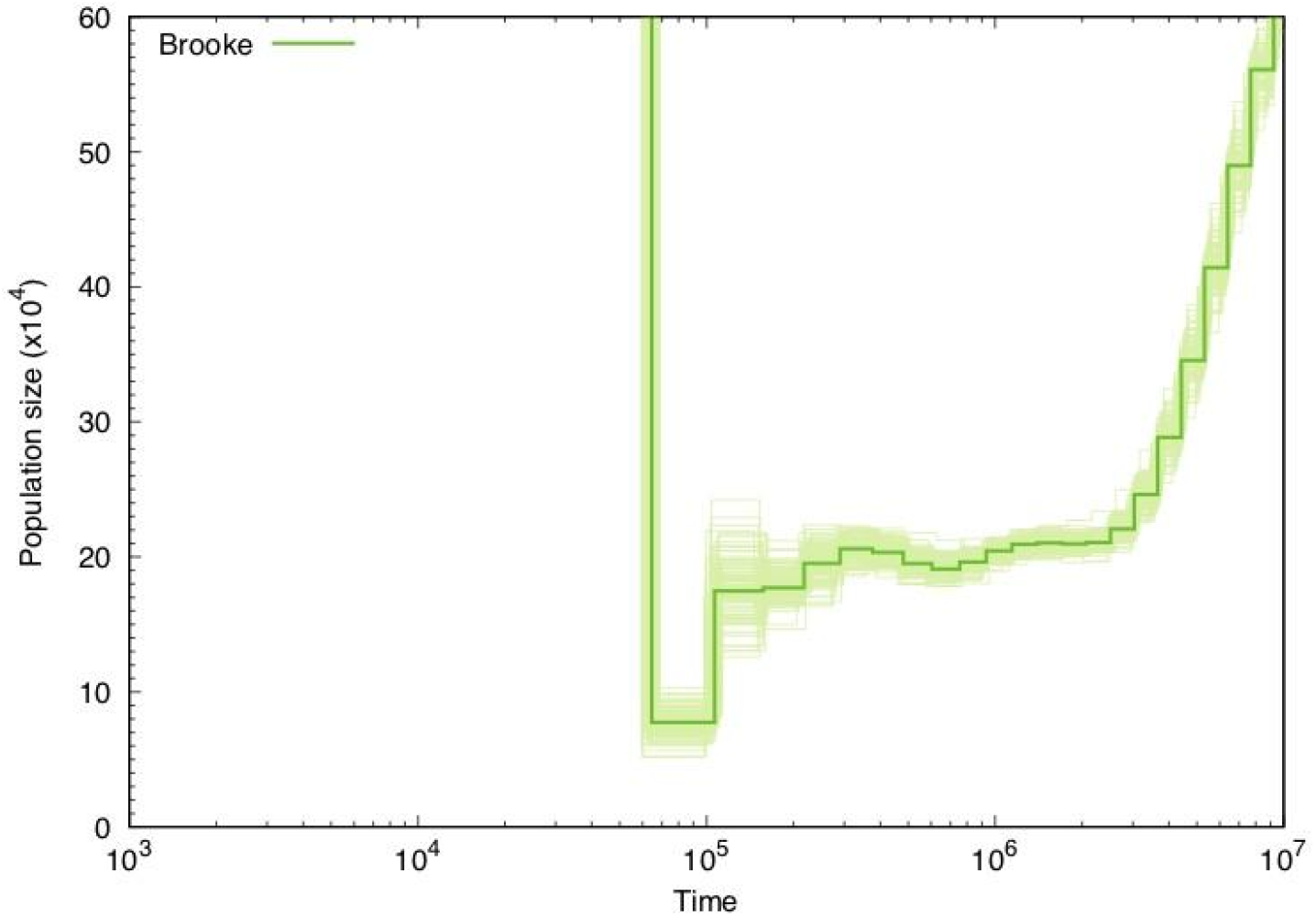
Bootstrap PSMC plot of lion sequenced in this study (‘Brooke’).

**Figure S5:**
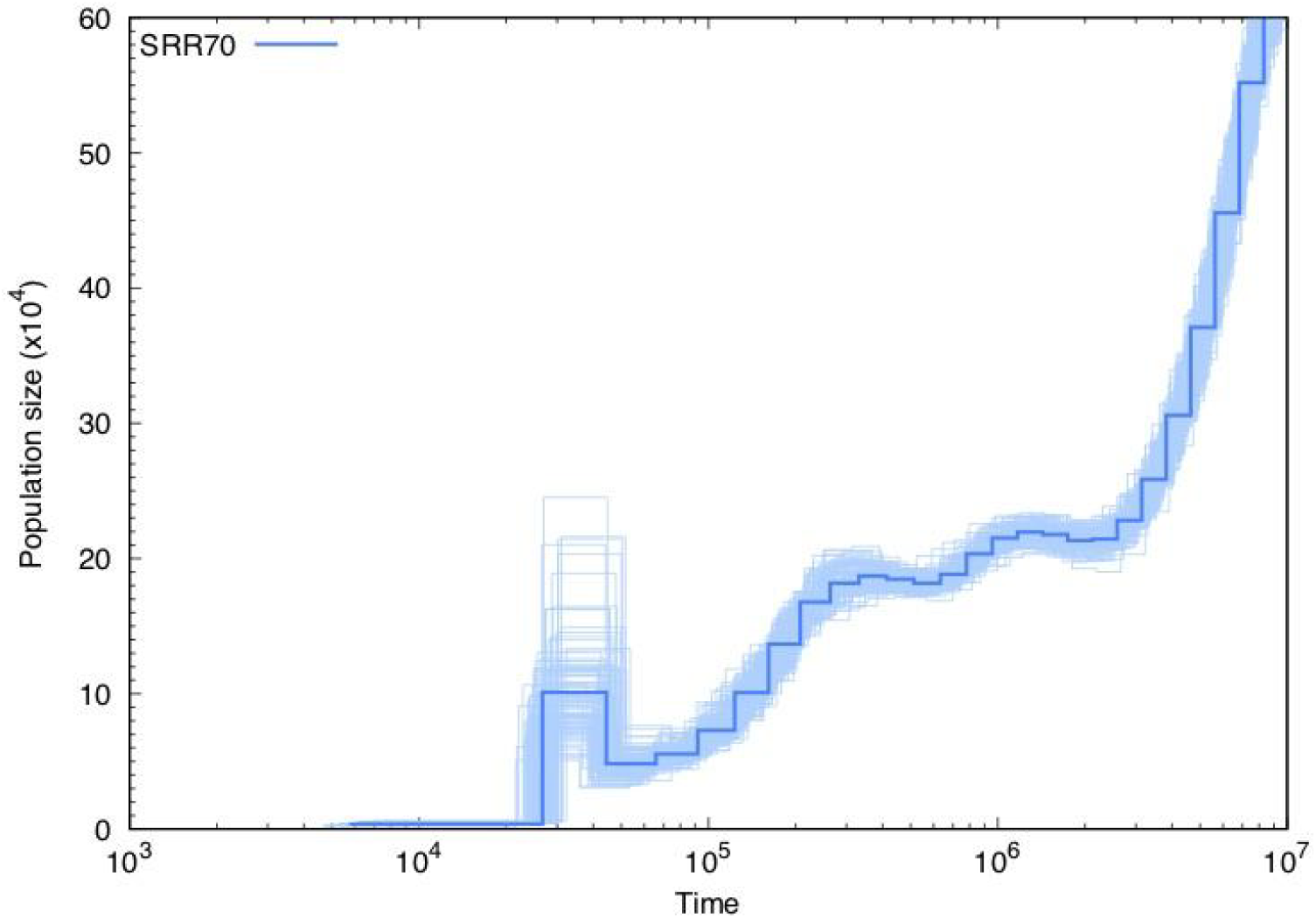
Bootstrap PSMC plot of white lion (Cho et al. 2013).

**Figure S6:**
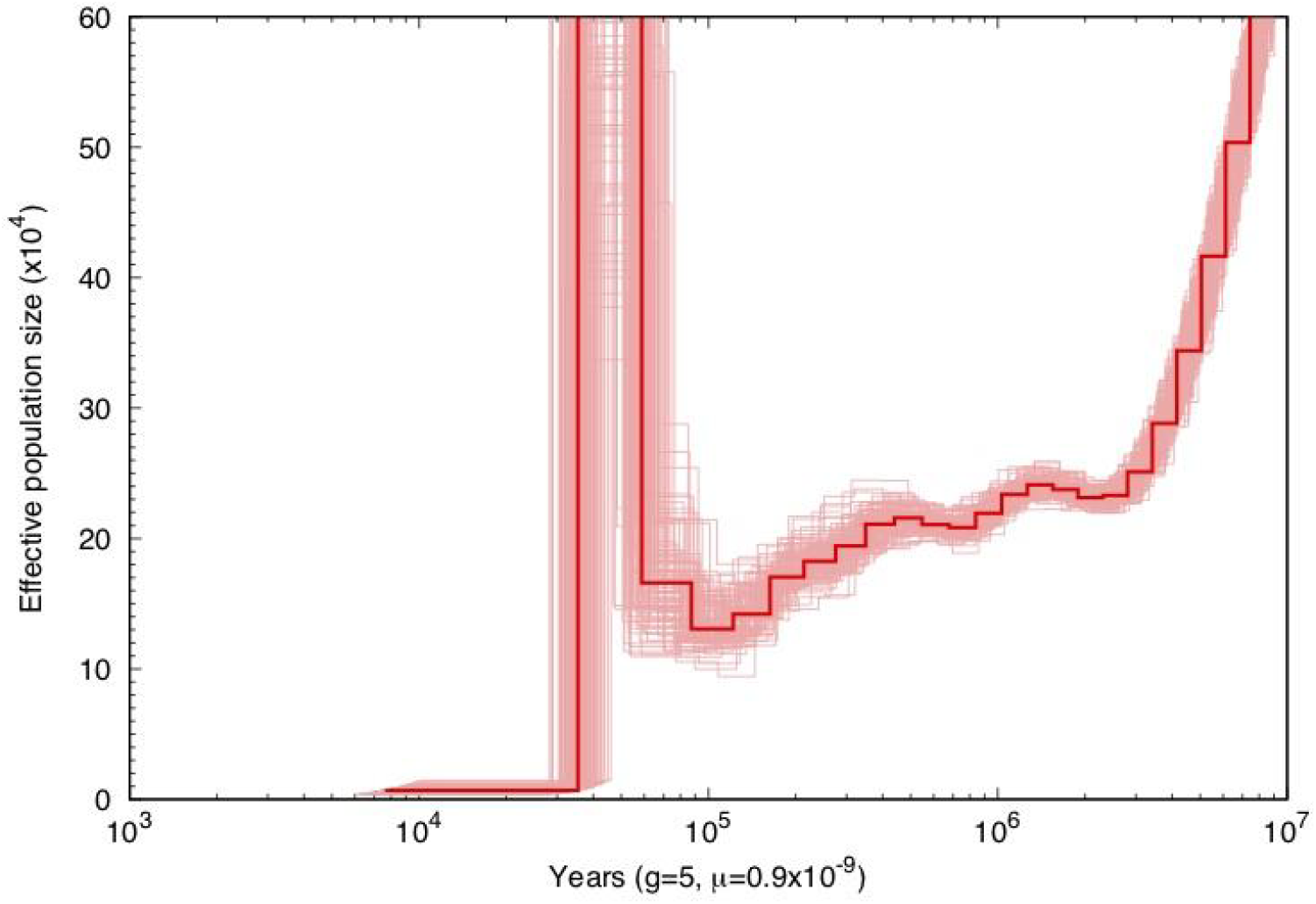
Bootstrap PSMC plot of tawny lion (Cho et al. 2013).

## References

1. Barnett R, Yamaguchi N, Barnes I, Cooper A. The origin, current diversity and future conservation of the modern lion (Panthera leo). Proc Biol Sci. royalsocietypublishing.org; 2006;273: 2119–2125.

2. Barnett R, Shapiro B, Barnes I, Ho SYW, Burger J, Yamaguchi N, et al. Phylogeography of lions (Panthera leo ssp.) reveals three distinct taxa and a late Pleistocene reduction in genetic diversity. Mol Ecol. Wiley Online Library; 2009;18: 1668–1677.

3. Wildt DE, Bush M, Goodrowe KL, Packer C, Pusey AE, Brown JL, et al. Reproductive and genetic consequences of founding isolated lion populations. Nature. nature.com; 1987;329: 328–331.

4. Ramanathan A, Malik PK, Prasad G. Seroepizootiological survey for selected viral infections in captive Asiatic lions (Panthera leo persica) from western India. J Zoo Wildl Med. BioOne; 2007;38: 400–408.

5. Bertola LD, Van Hooft WF, Vrieling K, Uit de Weerd DR, York DS, Bauer H, et al. Genetic diversity, evolutionary history and implications for conservation of the lion (Panthera leo) in West and Central Africa. J Biogeogr. Wiley Online Library; 2011;38: 1356–1367.

6. Bertola LD, Jongbloed H, van der Gaag KJ, de Knijff P, Yamaguchi N, Hooghiemstra H, et al. Phylogeographic Patterns in Africa and High Resolution Delineation of Genetic Clades in the Lion (Panthera leo). Sci Rep. nature.com; 2016;6: 30807.

7. Tensen L, Groom RJ, Khuzwayo J, Jansen van Vuuren B. The genetic tale of a recovering lion population (Panthera leo) in the Savé Valley region (Zimbabwe): A better understanding of the history and managing the future. PLoS One. journals.plos.org; 2018;13: e0190369.

8. Spong G, Stone J, Creel S, Björklund M. Genetic structure of lions (Panthera leo L.) in the Selous Game Reserve: implications for the evolution of sociality. J Evol Biol. Wiley Online Library; 2002;15: 945–953.

9. Munson L, Terio KA, Kock R, Mlengeya T, Roelke ME, Dubovi E, et al. Climate extremes promote fatal co-infections during canine distemper epidemics in African lions. PLoS One. journals.plos.org; 2008;3: e2545.

10. Munson L, Brown JL, Bush M, Packer C, Janssen D, Reiziss SM, et al. Genetic diversity affects testicular morphology in free-ranging lions (Panthera leo) of the Serengeti Plains and Ngorongoro Crater. J Reprod Fertil. rep.bioscientifica.com; 1996;108: 11–15.

11. Dubach JM, Briggs MB, White PA, Ament BA, Patterson BD. Genetic perspectives on “Lion Conservation Units” in Eastern and Southern Africa. Conserv Genet. Springer; 2013;14: 741–755.

12. Dubach J, Patterson BD, Briggs MB, Venzke K, Flamand J, Stander P, et al. Molecular genetic variation across the southern and eastern geographic ranges of the African lion, Panthera leo. Conserv Genet. Springer; 2005;6: 15–24.

13. Antunes A, Troyer JL, Roelke ME, Pecon-Slattery J, Packer C, Winterbach C, et al. The evolutionary dynamics of the lion Panthera leo revealed by host and viral population genomics. PLoS Genet. journals.plos.org; 2008;4: e1000251.

14. Singh A, Shailaja K, Gaur A, Singh L. Development and characterization of novel microsatellite markers in the Asiatic lion (Panthera leo persica). Mol Ecol Notes. Wiley Online Library; 2002;2: 542–543.

15. Bruche S, Gusset M, Lippold S, Barnett R, Eulenberger K, Junhold J, et al. A genetically distinct lion (Panthera leo) population from Ethiopia. Eur J Wildl Res. Springer; 2013;59: 215–225.

16. Miller SM, Harper CK, Bloomer P, Hofmeyr J, Funston PJ. Evaluation of microsatellite markers for populations studies and forensic identification of African lions (Panthera leo). J Hered. academic.oup.com; 2014;105: 762–772.

17. Gaur A, Shailaja K, Singh A, Arunabala V, Satyarebala B, Singh L. Twenty polymorphic microsatellite markers in the Asiatic lion (Panthera leo persica). Conserv Genet. Springer; 2006;7: 1005–1008.

18. Smitz N, Jouvenet O, Ambwene Ligate F, Crosmary W-G, Ikanda D, Chardonnet P, et al. A genome-wide data assessment of the African lion (Panthera leo) population genetic structure and diversity in Tanzania. PLoS One. journals.plos.org; 2018;13: e0205395.

19. Wurster-Hill DH, Gray CW. The interrelationships of chromosome banding patterns in procyonids, viverrids, and felids. Cytogenet Cell Genet. karger.com; 1975;15: 306–331.

20. Wurster-Hill DH, Centerwall WR. The interrelationships of chromosome banding patterns in canids, mustelids, hyena, and felids. Cytogenet Cell Genet. karger.com; 1982;34: 178–192.

21. Gopalakrishnan S, Samaniego Castruita JA, Sinding M-HS, Kuderna LFK, Räikkönen J, Petersen B, et al. The wolf reference genome sequence (Canis lupus lupus) and its implications for Canis spp. population genomics. BMC Genomics. bmcgenomics.biomedcentral.com; 2017;18: 495.

22. Brandt DYC, Aguiar VRC, Bitarello BD, Nunes K, Goudet J, Meyer D. Mapping Bias Overestimates Reference Allele Frequencies at the HLA Genes in the 1000 Genomes Project Phase I Data. G3. g3journal.org; 2015;5: 931–941.

23. Cueto M, Camarós E, Castaños P, Ontañón R, Arias P. Under the Skin of a Lion: Unique Evidence of Upper Paleolithic Exploitation and Use of Cave Lion (Panthera spelaea) from the Lower Gallery of La Garma (Spain). PLoS One. journals.plos.org; 2016;11: e0163591.

24. Goldman MJ, de Pinho JR, Perry J. Beyond ritual and economics: Maasai lion hunting and conservation politics. Oryx. Cambridge University Press; 2013;47: 490–500.

25. Hazzah L, Bath A, Dolrenry S, Dickman A, Frank L. From Attitudes to Actions: Predictors of Lion Killing by Maasai Warriors. PLoS One. journals.plos.org; 2017;12: e0170796.

26. Dolrenry S, Hazzah L, Frank LG. Conservation and monitoring of a persecuted African lion population by Maasai warriors. Conserv Biol. Wiley Online Library; 2016;30: 467–475.

27. Loveridge AJ, Searle AW, Murindagomo F, Macdonald DW. The impact of sport-hunting on the population dynamics of an African lion population in a protected area. Biol Conserv. Elsevier; 2007;134: 548–558.

28. Packer C, Brink H, Kissui BM, Maliti H, Kushnir H, Caro T. Effects of trophy hunting on lion and leopard populations in Tanzania. Conserv Biol. Wiley Online Library; 2011;25: 142–153.

29. Nelson F, Lindsey P, Balme G. Trophy hunting and lion conservation: a question of governance? Oryx. Cambridge University Press; 2013;47: 501–509.

30. Lindsey PA, Roulet PA, Romañach SS. Economic and conservation significance of the trophy hunting industry in sub-Saharan Africa. Biol Conserv. Elsevier; 2007;134: 455–469.

31. Williams VL, Loveridge AJ, Newton DJ, Macdonald DW. A roaring trade? The legal trade in Panthera leo bones from Africa to East-Southeast Asia. PLoS One. journals.plos.org; 2017;12: e0185996.

32. Trinkel M, Ferguson N, Reid A, Reid C, Somers M, Turelli L, et al. Translocating lions into an inbred lion population in the Hluhluwe-iMfolozi Park, South Africa. Anim Conserv. Wiley Online Library; 2008;11: 138–143.

33. Hunter LTB, Pretorius K, Carlisle LC, Rickelton M, Walker C, Slotow R, et al. Restoring lions Panthera leo to northern KwaZulu-Natal, South Africa: short-term biological and technical success but equivocal long-term conservation. Oryx. Cambridge University Press; 2007;41: 196–204.

34. Kyriazis CC, Wayne RK, Lohmueller KE. High genetic diversity can contribute to extinction in small populations [Internet]. bioRxiv. 2019. p. 678524. doi:10.1101/678524

35. Wright B, Farquharson KA, McLennan EA, Belov K, Hogg CJ, Grueber CE. From reference genomes to population genomics: comparing three reference-aligned reduced-representation sequencing pipelines in two wildlife species. BMC Genomics. bmcgenomics.biomedcentral.com; 2019;20: 453.

36. McCartney-Melstad E, Vu JK, Shaffer HB. Genomic data recover previously undetectable fragmentation effects in an endangered amphibian. Mol Ecol. Wiley Online Library; 2018;27: 4430–4443.

37. Parejo M, Henriques D, Pinto MA, Soland-Reckeweg G, Neuditschko M. Empirical comparison of microsatellite and SNP markers to estimate introgression in Apis mellifera mellifera. J Apic Res. Taylor & Francis; 2018;57: 504–506.

38. Mitra S, Sreenivas A, Sowpati DT, Kumar AS, Awasthi G, Kumar M, et al. De novo assembly and annotation of Asiatic lion (Panthera leo persica) genome [Internet]. bioRxiv. 2019. p. 549790. doi:10.1101/549790

39. Bagatharia SB, Joshi MN, Pandya RV, Pandit AS, Patel RP, Desai SM, et al. Complete mitogenome of Asiatic lion resolves phylogenetic status within Panthera. BMC Genomics. bmcgenomics.biomedcentral.com; 2013;14: 572.

40. Mohr DW, Naguib A, Weisenfeld N, Kumar V, Shah P, Church DM, et al. Improved de novo Genome Assembly: Linked-Read Sequencing Combined with Optical Mapping Produce a High Quality Mammalian Genome at Relatively Low Cost [Internet]. bioRxiv. 2017. p. 128348. doi:10.1101/128348

41. Putnam NH, O’Connell BL, Stites JC, Rice BJ, Blanchette M, Calef R, et al. Chromosome-scale shotgun assembly using an in vitro method for long-range linkage. Genome Res. genome.cshlp.org; 2016;26: 342–350.

42. Li H, Durbin R. Fast and accurate short read alignment with Burrows–Wheeler transform. Bioinformatics. Oxford University Press; 2009;25: 1754–1760.

43. Edgar RC. MUSCLE: multiple sequence alignment with high accuracy and high throughput. Nucleic Acids Res. 2004;32: 1792–1797.

44. Rice P, Longden I, Bleasby A. EMBOSS: the European Molecular Biology Open Software Suite. Trends Genet. 2000;16: 276–277.

45. Bradnam KR, Fass JN, Alexandrov A, Baranay P, Bechner M, Birol I, et al. Assemblathon 2: evaluating de novo methods of genome assembly in three vertebrate species. Gigascience. 2013;2: 10.

46. Simão FA, Waterhouse RM, Ioannidis P, Kriventseva EV, Zdobnov EM. BUSCO: assessing genome assembly and annotation completeness with single-copy orthologs. Bioinformatics. 2015;31: 3210–3212.

47. Katoh K, Standley DM. MAFFT multiple sequence alignment software version 7: improvements in performance and usability. Mol Biol Evol. 2013;30: 772–780.

48. Stamatakis A. RAxML version 8: a tool for phylogenetic analysis and post-analysis of large phylogenies. Bioinformatics. 2014;30: 1312–1313.

49. Mirarab S, Reaz R, Bayzid MS, Zimmermann T, Swenson MS, Warnow T. ASTRAL: genome-scale coalescent-based species tree estimation. Bioinformatics. 2014;30: i541–8.

50. Jurka J, Kapitonov VV, Pavlicek A, Klonowski P, Kohany O, Walichiewicz J. Repbase Update, a database of eukaryotic repetitive elements. Cytogenet Genome Res. 2005;110: 462–467.

51. Smit AFA, Hubley R, Green P. RepeatMasker. 1996.

52. Smit AFA, Hubley R. RepeatModeler Open-1.0. Available fom http://www.repeatmasker.org. 2008;

53. Tang H, Krishnakuar V, Li J. jcvi: JCVI utility libraries. Zenodo (doi: 105281/zenodo31631). 2015;

54. Kiełbasa SM, Wan R, Sato K, Horton P, Frith MC. Adaptive seeds tame genomic sequence comparison. Genome Res. genome.cshlp.org; 2011;21: 487–493.

55. Korneliussen TS, Albrechtsen A, Nielsen R. ANGSD: Analysis of Next Generation Sequencing Data. BMC Bioinformatics. 2014;15: 356.

56. Cho YS, Hu L, Hou H, Lee H, Xu J, Kwon S, et al. The tiger genome and comparative analysis with lion and snow leopard genomes. Nat Commun. 2013;4: 2433.

57. Dudchenko O, Shamim MS, Batra SS, Durand NC, Musial NT, Mostofa R, et al. The Juicebox Assembly Tools module facilitates de novo assembly of mammalian genomes with chromosome-length scaffolds for under $1000 [Internet]. bioRxiv. 2018. p. 254797. doi:10.1101/254797

58. Dudchenko O, Batra SS, Omer AD, Nyquist SK, Hoeger M, Durand NC, et al. De novo assembly of the Aedes aegypti genome using Hi-C yields chromosome-length scaffolds. Science. science.sciencemag.org; 2017;356: 92–95.

59. Armstrong E, Khan A, Taylor RW, Gouy A, Greenbaum G, Thiéry A, et al. Recent evolutionary history of tigers highlights contrasting roles of genetic drift and selection [Internet]. bioRxiv. 2019. p. 696146. doi:10.1101/696146

60. Wysoker A, Fennell T, Ruan J, Homer N, Marth G, Abecasis G, et al. The Sequence alignment/map (SAM) format and SAMtools. Bioinformatics. 2009;25: 2078–2079.

61. Dobrynin P, Liu S, Tamazian G, Xiong Z, Yurchenko AA, Krasheninnikova K, et al. Genomic legacy of the African cheetah, Acinonyx jubatus. Genome Biol. 2015;16: 277.

62. Kim S, Cho YS, Kim H-M, Chung O, Kim H, Jho S, et al. Comparison of carnivore, omnivore, and herbivore mammalian genomes with a new leopard assembly. Genome Biol. 2016;17: 211.

63. Abascal F, Corvelo A, Cruz F, Villanueva-Cañas JL, Vlasova A, Marcet-Houben M, et al. Extreme genomic erosion after recurrent demographic bottlenecks in the highly endangered Iberian lynx. Genome Biol. 2016;17: 251.

64. Saremi NF, Supple MA, Byrne A, Cahill JA, Coutinho LL, Dalén L, et al. Mountain lion genomes provide insights into genetic rescue of inbred populations [Internet]. bioRxiv. 2018. p. 482315. doi:10.1101/482315

65. Davis BW, Li G, Murphy WJ. Supermatrix and species tree methods resolve phylogenetic relationships within the big cats, Panthera (Carnivora: Felidae). Mol Phylogenet Evol. Elsevier; 2010;56: 64–76.

66. Johnson WE, Eizirik E, Pecon-Slattery J, Murphy WJ, Antunes A, Teeling E, et al. The late Miocene radiation of modern Felidae: a genetic assessment. Science. science.sciencemag.org; 2006;311: 73–77.

67. Robinson JA, Räikkönen J, Vucetich LM, Vucetich JA, Peterson RO, Lohmueller KE, et al. Genomic signatures of extensive inbreeding in Isle Royale wolves, a population on the threshold of extinction. Sci Adv. advances.sciencemag.org; 2019;5: eaau0757.

68. Ripple WJ, Van Valkenburgh B. Linking Top-down Forces to the Pleistocene Megafaunal Extinctions. Bioscience. Narnia; 2010;60: 516–526.

69. Lister AM, Stuart AJ. The impact of climate change on large mammal distribution and extinction: Evidence from the last glacial/interglacial transition. C R Geosci. Elsevier; 2008;340: 615–620.

70. Nadachowska-Brzyska K, Burri R, Smeds L, Ellegren H. PSMC analysis of effective population sizes in molecular ecology and its application to black-and-white Ficedula flycatchers. Mol Ecol. Wiley Online Library; 2016;25: 1058–1072.

71. Armstrong EE, Taylor RW, Prost S, Blinston P, van der Meer E, Madzikanda H, et al. Cost-effective assembly of the African wild dog (Lycaon pictus) genome using linked reads. Gigascience. academic.oup.com; 2019;8. doi:10.1093/gigascience/giy124

72. 10K Community of Scientists G. Genome 10K: A Proposal to Obtain Whole-Genome Sequence for 10 000 Vertebrate Species. J Hered. Narnia; 2009;100: 659–674.

73. Koepfli K-P, Paten B, of Scientists G 10k C, O’Brien SJ. The Genome 10K Project: a way forward. Annu Rev Anim Biosci. Annual Reviews; 2015;3: 57–111.

74. Consortium I. The i5K Initiative: Advancing Arthropod Genomics for Knowledge, Human Health, Agriculture, and the Environment. J Hered. Narnia; 2013;104: 595–600.

75. Zhang G, Rahbek C, Graves GR, Lei F, Jarvis ED, Gilbert MTP. Genomics: Bird sequencing project takes off. Nature. nature.com; 2015;522: 34.

76. Packer C, Pusey AE, Rowley H, Gilbert DA, Martenson J, O’brien SJ. Case Study of a Population Bottleneck: Lions of the Ngorongoro Crater. Conserv Biol. Wiley Online Library; 1991;5: 219–230.

77. Packer C. The African Lion: A Long History of Interdisciplinary Research. Frontiers in Ecology and Evolution. frontiersin.org; 2019;7: 259.

